# Quantifying the influence of biophysical factors in shaping brain communication through remnant functional networks

**DOI:** 10.1101/2024.09.11.612501

**Authors:** Johan Nakuci, Javier Garcia, Kanika Bansal

## Abstract

Functional connectivity (FC) reflects brain-wide communication essential for cognition, yet the role of underlying biophysical factors in shaping FC remains unclear. We quantify the influence of physical factors—structural connectivity (SC) and Euclidean distance (DC), which capture anatomical wiring and regional distance—and molecular factors—gene expression similarity (GC) and neuroreceptor congruence (RC), representing neurobiological similarity—on resting-state FC. We assess how these factors impact graph-theoretic and gradient features, capturing pairwise and higher-order interactions. By generating *remnant functional networks* after selectively removing connections tied to specific factors, we show that molecular factors, particularly RC, dominate graph-theoretic features, while gradient features are shaped by a mix of molecular and physical factors, especially GC and DC. SC has a surprisingly minor role. We also link FC alterations to biophysical factors in schizophrenia, bipolar disorder, and ADHD, with physical factors differentiating these groups. These insights are key for understanding FC across various applications, including task performance, development, and clinical conditions.

## Introduction

Communication among brain regions is fundamental to function. Functional connectivity (FC) provides a compact view of the fundamental features of brain-wide communication (Bassett and Sporns, 2017) essential for cognitive processes (Chadick and Gazzaley, 2011; Cole et al., 2016). However, how the features of FC and intrinsically linked regional brain communication emerge from the underlying biophysical constraints is poorly understood.

These features have been primarily investigated using graph-theoretic measures which quantify pairwise relationships. Graph-theoretic measures have identified components facilitating efficient information transfer (Avena-Koenigsberger et al., 2017), global integration and local segregation (Cohen and D’Esposito, 2016), and robustness to injury crucial for effective communication (Aerts et al., 2016). Additionally, recent work has shown that alongside pairwise relationships, higher-order interactions associated with the gradient of the FC, are also a fundamental feature of brain communication (Margulies et al., 2016).

Most studies and modeling frameworks have focused on understanding the role of anatomical wiring or structural connectivity (SC) in constraining FC, which provides a fundamental *physical* basis for brain communication (Honey et al., 2009). Nonetheless, other factors have been shown to shape brain-wide communication. Another physical factor is Euclidean distance (distance dependent connectivity or DC) which allows for easier and/or faster communication between regions that are in closer proximity than the ones at farther physical distance (Shinn et al., 2023). Brain communication is also fundamentally constrained and shaped by molecular relationships which reflect the similarity in the neurobiological make-up between regions. Previous studies have captured these relationships using similarity in genetic expression (GC) (Richiardi et al., 2015) and neuroreceptor density (RC) (Hansen et al., 2022a). Here, we consider these *biophysical factors* collectively to uncover the extent to which physical and molecular factors constrain and shape the fundamental features of FC.

This stands in contrast to the questions that have primarily been addressed thus far which have focused on whether *an individual* biophysical factor shapes and constrains FC while little attention has been focused on the question pertaining to the relative prominence between biophysical factors. The primary focus here is to determine the relative prominence of each of these biophysical factors in shaping and containing FC. We also investigate if different biophysical factors shape and constrain different aspects of the functional connectivity.

Addressing these questions is challenging and requires a method that is applicable across biophysical factors. Part of the challenge stems from the fact that biophysical factors are generated from different types of data and as a result reflect different relationships between brain regions. Existing comparative approaches in this direction correlate FC with biophysical properties, such as in Hansen et al., 2022a, but suffer from the limitation of assuming a linear relationship between FC and a biophysical factor which is not ideal considering that some biophysical factors such as structural connectivity and Euclidean distance do not exhibit a linear relationship with FC (Hansen et al., 2023). Given these challenges, a robust method is needed that can be applicable across multiple biophysical factors and does not make strong prior assumptions about the relationship between biophysical factors and FC.

Here, we developed a simple test to quantify the underlying influence of biophysical factors based on remnant functional networks (RFNs) to overcome these challenges. The first step in the framework is to derive a network for each of the biophysical factors that quantifies the pairwise relationship between brain regions. We then compute the alteration in organizational features of the FC network before and after the connections shared with a reference biophysical network are removed. We call these FC connectivity maps with removed connections *remnant functional networks* (RFNs, **Fig. 1**). Observed differences between the organizational features of the FC and various RFNs inform us on how a biophysical network shapes FC.

**Figure 1.**
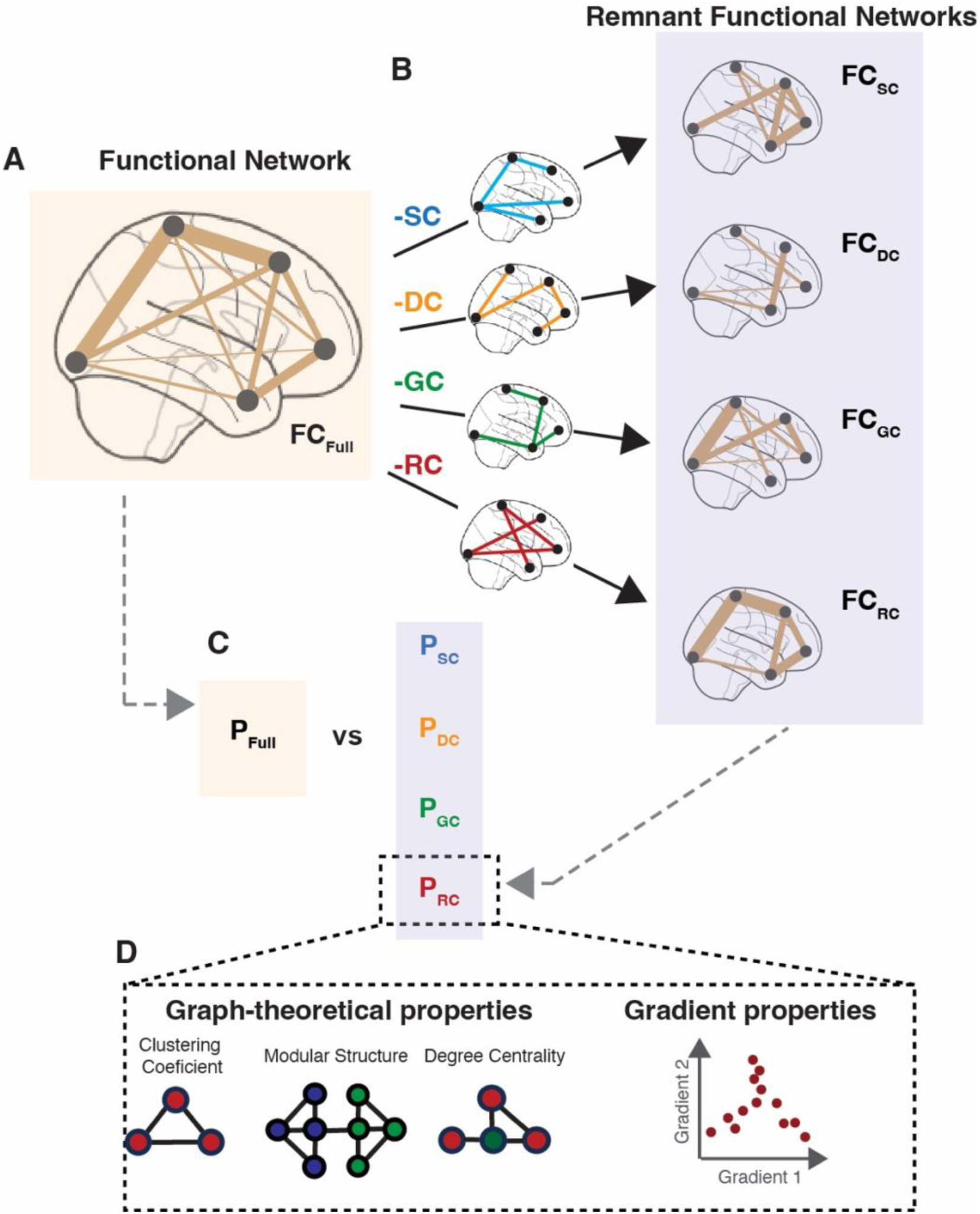
Schematic of the analytical test based on remnant functional networks. (A) Fully connected weighted functional connectivity network (FC_Full_). The width of the lines represents the strength of the connection. (B) We derive remnant functional networks (RFNs) by removing the connections in FC_Full_ that share a direct connection in an underlying biophysical network. These biophysical networks are structural connectivity (SC), Euclidean distance (DC), similarity in genetic expression (GC), and neuroreceptor congruence (RC) between brain regions; and corresponding RFNs are denoted by FC_SC_, FC_DC_, FC_GC_, and FC_RC_ respectively. (C) To assess the role of these biophysical networks in shaping the organization of the brain’s functional connectivity, we compare the features (P) of FC_SC_, FC_DC_, FC_GC_, and FC_RC_ (i.e., P_SC_, P_DC_, P_GC_, and P_RC_) with the features of FC_Full_ (i.e., P_Full_). (D) The features we employ comprise a comprehensive set of graph-theoretical and gradient features.

With the RFN, we can answer which of the biophysical factors plays the prominent (i.e. strongest) role. Additionally, we can determine which biophysical factors shape and constrain various functional properties – graph-theoretic and gradients. Moreover, the advantage of RFN framework is that it makes comparison across biophysical factors possible despite the inherent differences among the factors. Additionally, RFNs do not require any *a priori* assumptions about a linear relationship between FC and biophysical factors.

We present our main findings with functional connectivity and RFNs obtained from the Human Connectome Project (HCP) (Van Essen et al., 2013), and also corroborate them in another publicly available dataset UCLA Consortium for Neuropsychiatric Phenomics Dataset (LA5c) (Poldrack et al., 2016). Additionally, we complement our analytical test with a modeling analysis that predicts functional connectivity from each of the biophysical networks. Lastly, we extend the framework to understand disease specific alterations, particularly neuropsychiatric disorders where deviations in connectivity patterns are likely subtle, to identify the biophysical networks contributing to cognitive disturbances.

## Results

### Reference functional connectivity and biophysical networks

The analysis aims to quantify the influence different biophysical networks have in constraining and shaping the foundational organization of brain functional connectivity estimated from fMRI derived BOLD signal. To accomplish this, we first estimated the functional connectivity for an individual subject from the Human Connectome Project (HCP, N=48), using the resting-state fMRI signal from 200 brain regions derived from the Schaefer atlas, and computed the Pearson correlation between pairs of brain regions (Schaefer et al., 2018). We then average FC networks across individuals to derive the reference functional connectivity – FC_Full_ (**Fig. 1A and Fig. 2A**).

**Figure 2.**
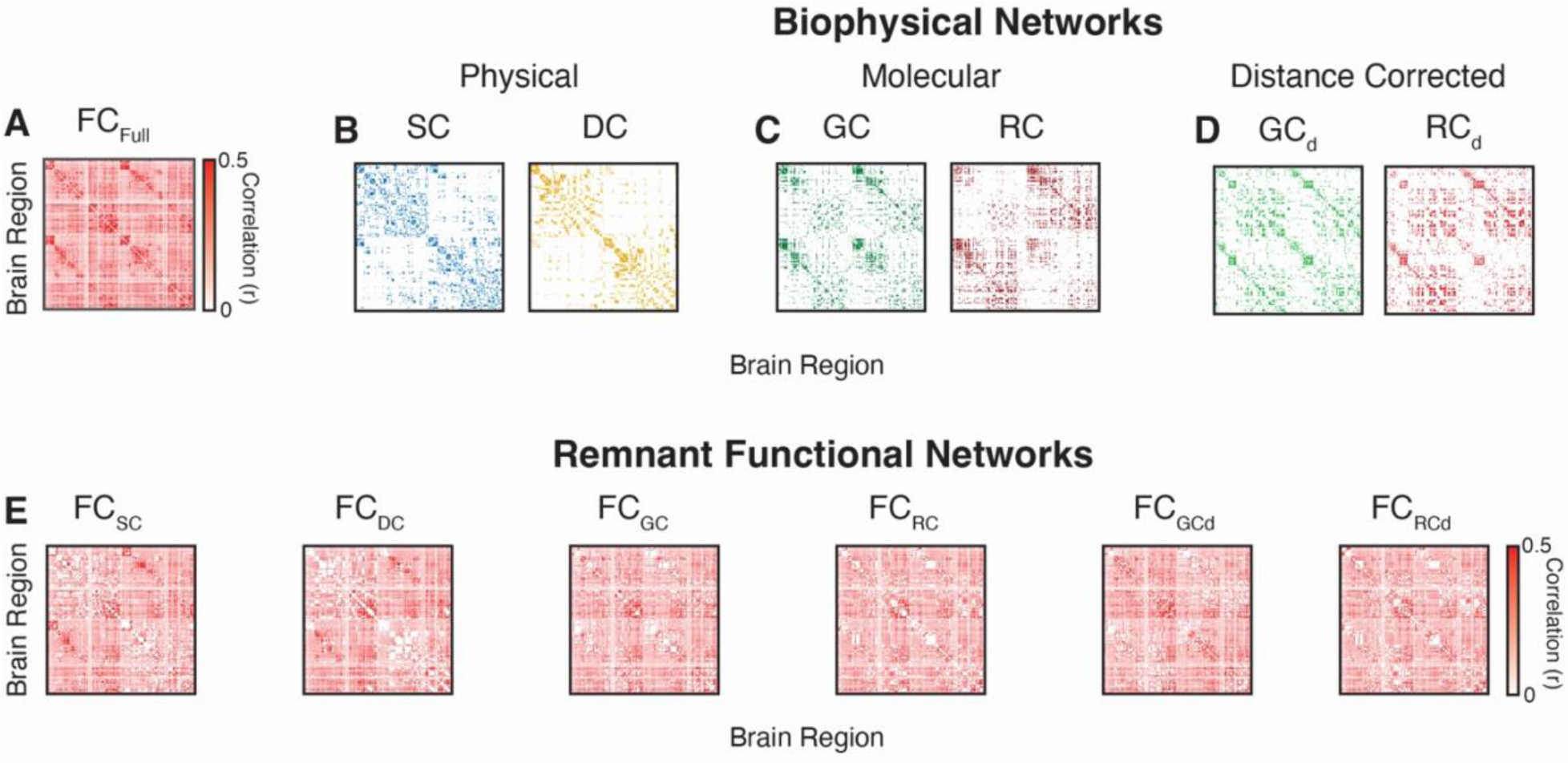
Connectivity patterns of group-level FC, SC, DC, GC and RC networks. (A) Functional brain connectivity (FC_Full_) between 200 brain regions derived from the Schaefer brain atlas. (B) structural connectivity (SC, *left*) and Euclidean distance (DC, *right*), networks reflecting physical constraints between brain regions. (C) Similarity in gene expression (GC, *left*) and neuroreceptor congruence (RC), networks reflecting molecular relationships between brain regions. (D) Distance corrected gene expression (GC_ED_, *left*) and neuroreceptor congruence (RC_ED_, *left*). (E) Remnant functional networks derived from each of the biophysical networks – FC_SC_, FC_DC_, FC_GC_, FC_RC_, FC_GCd_, and FC_RCd_.

As described in **Fig. 1**, our test used FC_Full_ to construct remnant functional networks (RFNs) by removing edges shared with a given biophysical network (**Fig. 1B**). Specifically, to create an RFN from a biophysical network, connections in FC that also had a corresponding connection in the reference biophysical network (e.g., SC, DC, GC or RC) were removed (**Fig. 1B)**. When the reference network was SC, procedurally this entailed setting all connections in the FC_Full_ network to zero that also have structural connections; this produces a weighted functional network with remnant connections to structural connectivity (FC_SC_; **Fig, 1C**). This procedure is repeated for DC, GC and RC-based networks resulting in weighted networks with only remnant connections, i.e., FC_DC_, FC_GC_, FC_RC_, respectively. Note that for Euclidean distance, the values were first converted from distance to proximity by subtracting the pairwise distance from the max distance between brain regions (see Methods for more details).

We then estimated several features of FC_Full_ and for each RFNs to quantify and compare the effect of a biophysical factor on a given network feature (**Fig. 1C**). We employed a variety of graph-theoretical as well as gradient features, capturing pair-wise as well as higher-order interactions, to gain a holistic understanding of the role of different biophysical factors (**Fig. 1D**).

Reference biophysical networks that reflect a diverse array of relationships between two brain regions were derived differently. We focus on two groups–physical and molecular networks– where physical networks reflect either the physical wiring or physical distance between two brain regions. To capture physical wiring, we derived group-level structural connectivity (SC) from white matter streamlines (**Fig. 2B**). To capture distance-based relationships, we derived distance dependent connectivity (DC) by computing Euclidean distance between the centroids of two corresponding brain regions in the Schaefer atlas (**Fig. 2B**).

For molecular relationships we focused on two distinct neurobiological factors pertaining to the regional similarities in gene expression (GC) and congruence in neuroreceptor density (RC). GC or the extent to which two brain regions express the same genes, was estimated from RNA microarray data using the Pearson correlation (**Fig. 2C**) (Hawrylycz et al., 2012). Similarly, RC or the extent to which two brain regions exhibit similar concentration of neuroreceptors was estimated from PET tracer imaging of 19 neuroreceptors and transporters maps using the Pearson correlations (**Fig. 2C**) (Hansen et al., 2022). Additionally, previous work has shown that the similarity in genetic expression and neuroreceptor congruence is dependent on the distance between brain regions (Shinn et al., 2023). To account for this dependence, we derived two additional distance corrected networks – GC_d_ and RC_d_ (**Fig. 2D**, see Methods for more details on molecular networks).

SC is a sparsely connected network due to the absence of a direct anatomical connection between several brain regions; other factors represent fully connected networks, meaning, a nonzero value connects all pairs of regions. This mismatch in the *density* of the networks does not allow for a comparison across different biophysical networks. Therefore, we matched the density of all the biophysical networks (DC, GC, RC) to SC ensuring that the same number of connections were removed when constructing the RFNs (see Methods). **Fig. 2E** shows the RFNs created by removing connections associated with different biophysical networks in FC_Full_. In the following sections we utilize RFNs to quantify the influence each of the biophysical networks has on the organization of functional brain connectivity.

### Majority of functional connections have an underlying biophysical substrate

Creating an RFN will result in many of the top functional connections, which are expected to strongly influence network organization, to be removed. However, the RFN may still retain strong functional connections without an underlying substrate. To investigate this, we thresholded the FC and biophysical networks preserving only the top certain fraction (%) of connections and assessed the extent of overlap between the thresholded FC and biophysical networks. We performed thresholding over a range of values preserving the top 5% to the top 50% of connections.

For the top 50% of strongest FC connections, 94.3% of these connections could be linked to at least one biophysical network (**Fig. 3A**). Even for the strongest top 5% of FC connections, 64.4% of these connections could be linked to at least structural connectivity or biophysical interaction. Moreover, the strongest functional connections consistently exhibited highest overlap with molecular factors – GC and RC (**Fig. 3B**). Taken together, these results suggest that underlying biophysical networks explain the majority (if not all) of the functional connections between brain regions.

**Figure 3.**
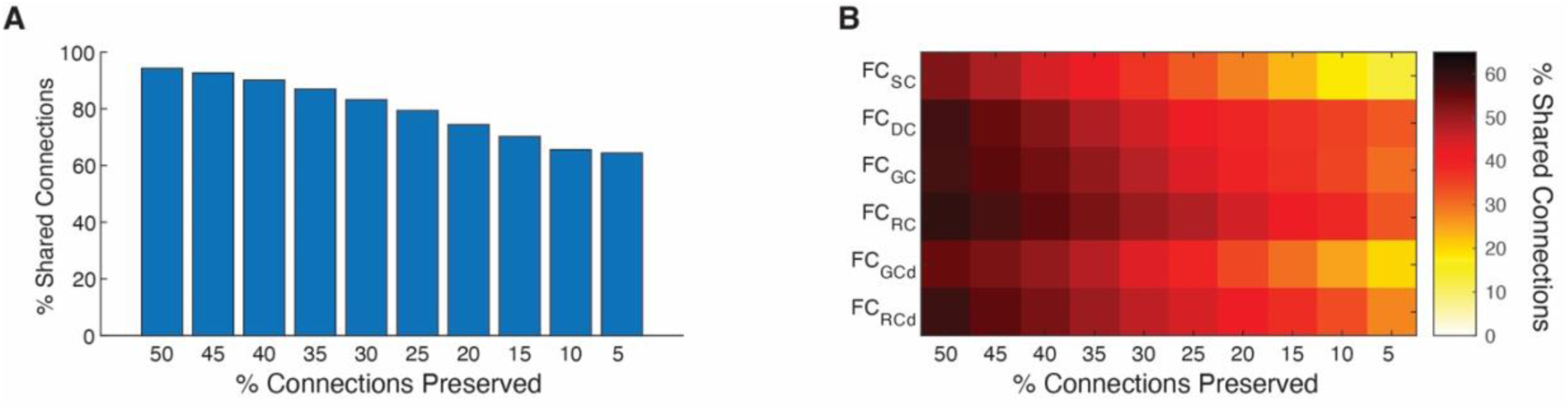
Majority of the strongest functional connections have an underlying biophysical substrate. (A) Strongest connections in FC can be linked to at least one biophysical network. The FC and biophysical network were thresholded to preserve the top X% of connections and the percentage of preserved connections in FC that were present in at least one biophysical network was estimated. FC and biophysical networks were thresholded over a range of values preserving the top 5% to the top 50% of connection. (B) Percentage of strongest connections in FC that can be linked to each biophysical network.

### Biophysical networks constraining and shaping graph-theoretical features of FC

Having established that the majority of strongest connection in FC can be linked to biophysical networks, we then focused on determining how biophysical networks shape pairwise relationships between brain regions by quantifying the changes in graph-theoretic features between FC_Full_ and RFNs. We leveraged a comprehensive set of features that have been identified to measure a diverse set of functions required for cognitive processing (Medaglia et al., 2015). These features include modularity, weighted degree, clustering coefficient, path length, spectral radius, eigenvector centrality, and synchronizability (see Methods for details).

Specifically, we calculated each of these features for the group-level functional connectivity network (P_Full_) and then assessed the percent changes in each of these features in the RFNs. Evaluating percent change allowed us to compare the relative shift in different features despite the differences in their magnitude. Note that to equate the density across biophysical networks, we estimated the average density of the SC network across subjects (d_ave_ = 16.07%) since it is the least dense compared to the other networks. Then each of the biophysical networks were thresholded so that the most prominent (i.e. strongest) 16.07% of connections were retained.

Comparing across graph-theoretic measures, removing shared connections with biophysical networks affected modularity the most (|Δ| = 24.27 ± 10.53% on an average; **Fig. 4A**). Modularity was followed by spectral radius, clustering coefficient, and weighted degree (∼ 20% change on average). Whereas eigenvector centrality showed the lowest change (< 1%). Across all graph-theoretic properties, the molecular networks, particularly RC, were the dominant factor. On average RC had the strongest impact (i.e. |Δ|) of 18.66 ± 10.70%, **Fig. 4B**). Even after correcting for the distance dependencies, RC_d_ had a higher impact than the other biophysical networks (|Δ| = 17.59 ± 9.75%). Moreover, the GC network closely followed the magnitude of RC in shaping FC. Interestingly, removing SC associated connections had the least effect on the organization of FC compared to all other biophysical networks.

**Figure 4.**
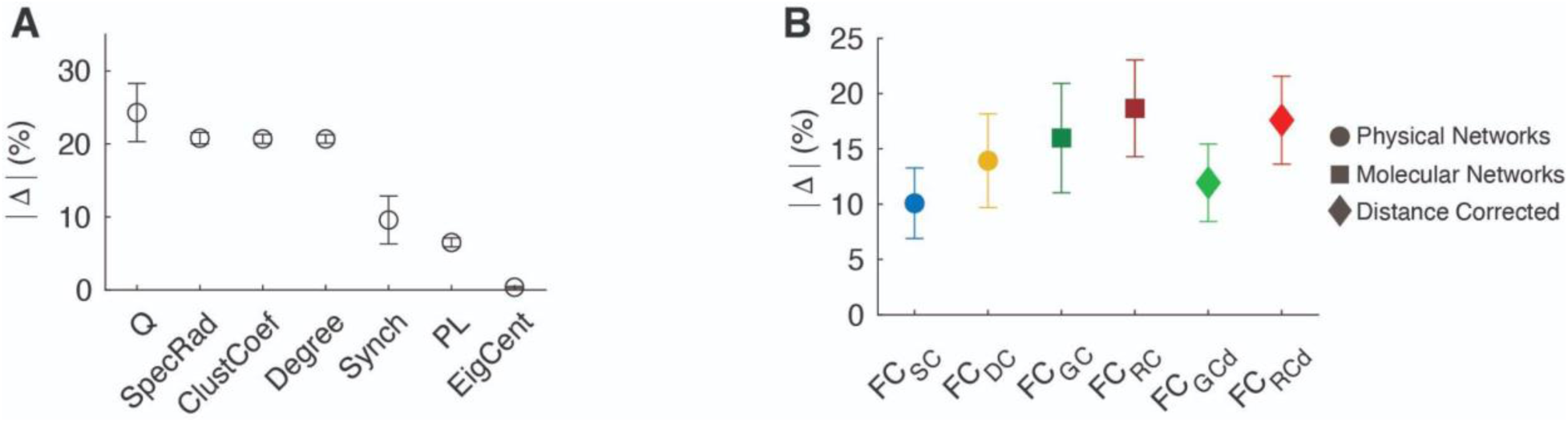
Molecular networks are the dominant factor constraining and shaping functional connectivity. (A) Average change in graph-theoretic metrics across biophysical networks. Weighted Degree (Degree), Clustering Coefficient (ClustCoef), Spectral Radius (SpecRad), Path Length (PL), Modularity (Q), Synchronizability (Synch) and Eigenvector Centrality (EigCent). (B) Average change for biophysical networks across graph-theoretic metrics, highlighting the dominance of molecular networks and RC in particular. For degree, clustering coefficient, and eigenvector centrality values were averaged across all brain regions.

Focusing on individual graph-theoretic properties, as can be observed in **Fig. 5A**, for weighted degree removing connections associated with each of the biophysical networks, resulted in a decrease in weighted degree in the RFNs compared to FC_Full_. The largest change occurred due to molecular factors – FC_RC_ (Δ = -22.07%) and FC_GC_ (Δ = -21.24%), followed by FC_DC_ (Δ = - 20.39%), and FC_SC_ (Δ = -17.99%). Moreover, we found that correcting for distance in RC and GC had a similar effect – FC_RCd_ (Δ = -22.08%), FC_GCd_ (Δ = -20.29%). Overall, neuroreceptor congruence emerged as a pivotal factor affecting weighted degree.

**Figure 5.**
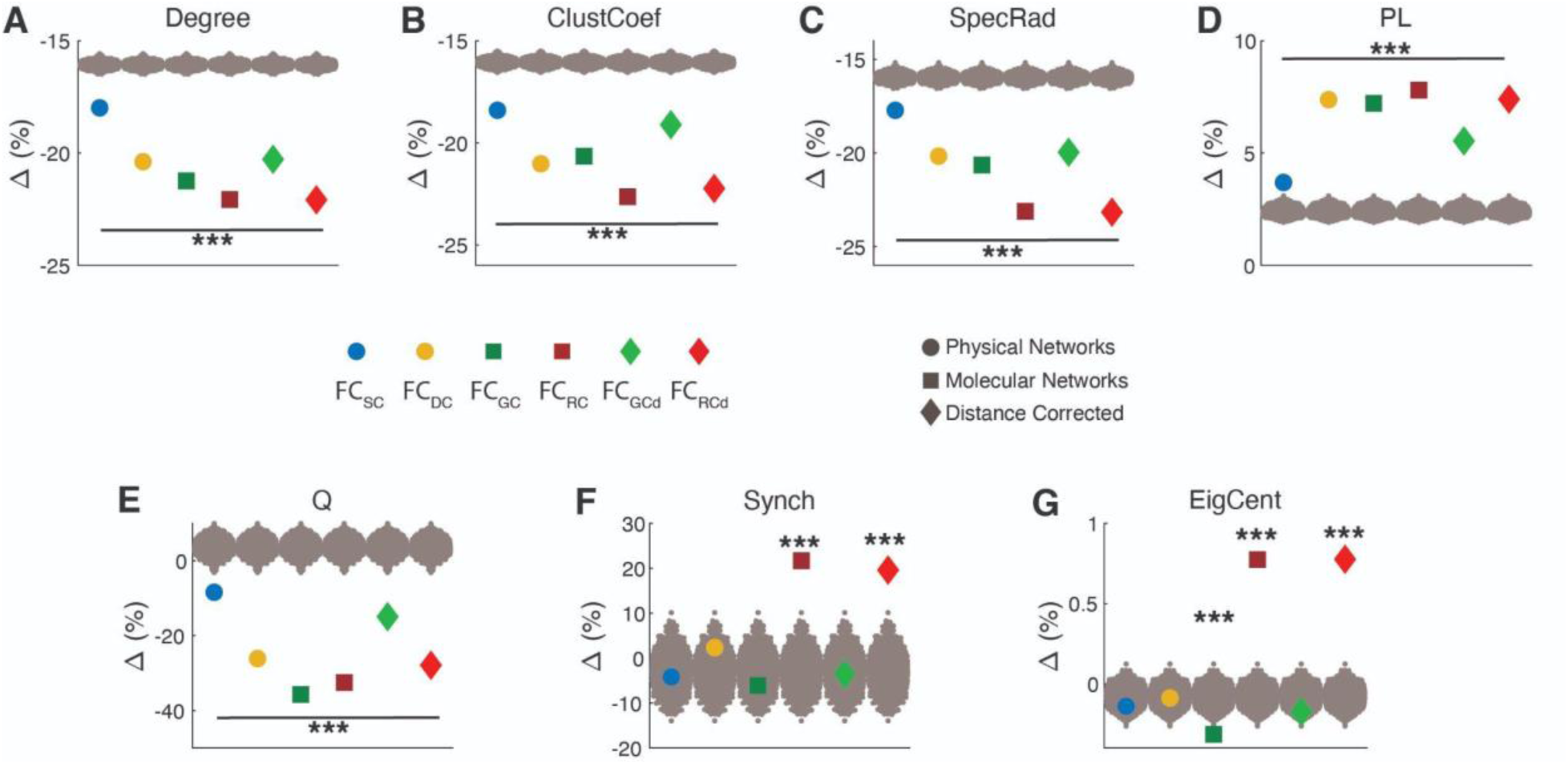
Biophysical networks shaping the graph-theoretical properties of functional brain connectivity. (A) Weighted Degree (Degree), (B) Clustering Coefficient (ClustCoef), (C) Spectral Radius (SpecRad), (D) Path Length (PL), (E) Modularity (Q), (F) Synchronizability (Synch) and (G) Eigenvector Centrality (EigCent). The effect of each biophysical network was quantified as a percent change (Δ) between FC_Full_ and RFNs for each graph-theoretic measure. Colored dots represent the values after removing SC, DC, GC, RC, GC_d_, and RC_d_ associated connections in the FC_Full_. For degree, clustering coefficient, and eigenvector centrality values were averaged across all brain regions. Gray dots represent the values after removing a random set of connections in the FC_Full_ (null model). FC_SC_: FC with SC associated connections removed; FC_DC_: FC with DC connections removed; FC_GC_: FC with GC connections removed; FC_RC_: FC with RC connections removed; FC_GCd_: FC with GC corrected for distance dependencies removed; FC_RCd_: FC with RC corrected for distance dependencies removed. Statistical testing assessed the deviation from the null model with *** denoting p-value < 0.001.

The weighted degree of the network, and other network properties, will inherently change when connections are removed, therefore, we tested if the observed changes were greater than what would be observed from removing random connections – null model. We randomly removed a set of connections from FC_Full_ that were equal to the number of connections removed by each of the biophysical networks and calculated the shift in graph-theoretic features. This procedure was repeated 1,000 times to generate a null distribution. The observed alterations in weighted degree induced by each of the biophysical networks were significantly greater than the null model (P_rand_ < 0.001).

Similarly, we observed the dominance of molecular factors on other network features. Particularly, RC-based networks (FC_RC_) exerted the dominant influence on average clustering coefficient (Δ = -22.64%, P_rand_ < 0.001; **Fig. 5B**), spectral radius (Δ = -23.16%, P_rand_ < 0.001; **Fig. 5C**) and path length (Δ = 7.80%, P_rand_ < 0.001; **Fig. 5D**). For modularity, GC-based network exerted the strongest influence (FC_GC_: Δ = -35.72%; P_rand_ < 0.001) closely followed by RC-based networks (FC_RC_: Δ = -32.56%; P_rand_ < 0.001); however, after correcting for distance, RC exerted the larger influence (FC_RCd_: Δ = -27.91%; P_rand_ < 0.001; **Fig. 5E**). Moreover, RC-based networks similarly exerted the largest influence on synchronizability – FC_RC_ (Δ = 21.67%, P_rand_ < 0.001; **Fig. 5F**), and eigenvector centrality – FC_RC_ (Δ = 0.78%, P_rand_ < 0.001; **Fig. 5G**).

In general, we observed a higher influence of molecular factors than physical factors, with structural connectivity showing the lowest percent change in all the graph-theoretic metrics tested. These results suggest that attempting to derive functional relationships from structural connections only has severe limitations and instead a greater emphasis should be placed on molecular factors (i.e. genetic expression and neuroreceptor congruence).

### Regional relationship between biophysical factors and graph-theoretic features of FC

Given the dominance of neuroreceptors congruence, we additionally probed the regional specificity of RC driven changes for graph-theoretic properties for which a value could be estimated for each brain region, i.e., weighted degree, eigenvector centrality, and clustering coefficient. For this analysis, percent change in these node-wise features was calculated at the region level and in **Figure 6** we show the observed changes as a brain-wide heat map. We found that the relationship between neuroreceptor congruence and these features was differentially distributed across the cortex. In particular, neuroreceptor congruence induced the largest shifts in the dorsolateral prefrontal cortex, motor and visual areas, whereas the medial prefrontal cortex and insula exhibited minor changes. It is worth noting, that while on average neuroreceptor congruence exhibited minimal effect on eigenvector centrality (**Fig. 5G**), this was a result of the differential relationship across the cortex (**Fig. 6C**). In fact, removing RC driven connectivity led to an increase in the eigenvector centrality of the medial prefrontal cortex, parietal cortex, insula and temporal lobe suggesting that neuroreceptor congruence functions to dampen the centrality profile of these brain regions. Moreover, the magnitude of change across brain regions for weighted degree and clustering coefficient is strongly associated with the strength of the feature indicating that the hub properties emerge from underlying neuroreceptor congruence (**Fig. 6D, Fig. S1**).

**Figure 6.**
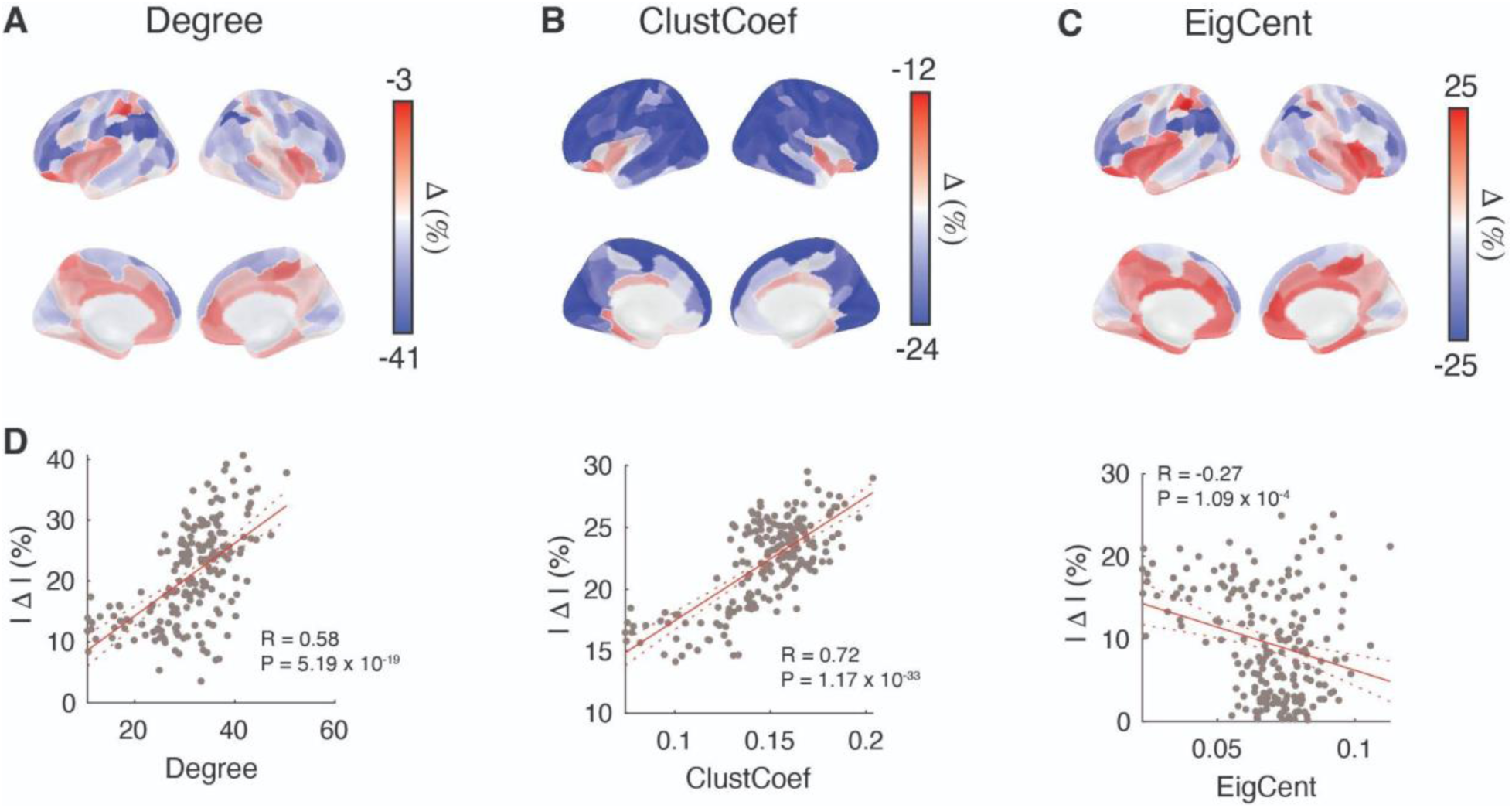
Neuroreceptor congruence dependent associations in functional connectivity are brain wide. Impact of neuroreceptor congruence on (A) weighted degree, (B) clustering coefficient, and (C) eigenvector centrality across the cortex. (D) Relationship between the regional network features and magnitude of change across brain regions.

### Biophysical networks constraining and shaping the gradient organization of FC

In addition to graph-theoretic features which focus on pairwise relationships between brain regions, gradient analysis can be used to infer higher-order relationships among brain regions. The gradient architecture of the cortex is one of the organizational features that is believed to anchor cortical functional hierarchy (Margulies et al., 2016). Therefore, we investigated how biophysical factors influence the gradient organization of FC to determine the underlying factors influencing and shaping functional hierarchy.

To unravel the hierarchical architecture within functional networks, Margulies et al., 2016 presented a framework based on topological data analysis to determine the components or *gradients* that describe the maximum variance in a network. Briefly, gradient analysis relies on computing the similarity in nodal connectivity patterns followed by dimensionality reduction to extract orthogonal components (gradients) that encode the dominant differences in nodes’ connectivity patterns. We leverage their framework to determine how the architecture of the primary gradients of RFNs differ from that of the brain’s functional connectivity in order to determine the role biophysical networks have in shaping the hierarchical architecture (see Methods for details).

In **Figure 7A** we show scatter plots of two primary connectivity gradients for different networks and color them based on the functional communities part of the Schaefer atlas. These primary gradients are the top two gradient components to describe the differences in nodal connectivity patterns. For FC_Full_, we replicate previously presented findings and relative organization of different communities. The observed organization places the default-mode network (DMN), somatomotor (SOM), and visual cortex (VIS) at opposite ends of the spectrum with the Gradient 2 largely separating the DMN from SOM.

**Figure 7.**
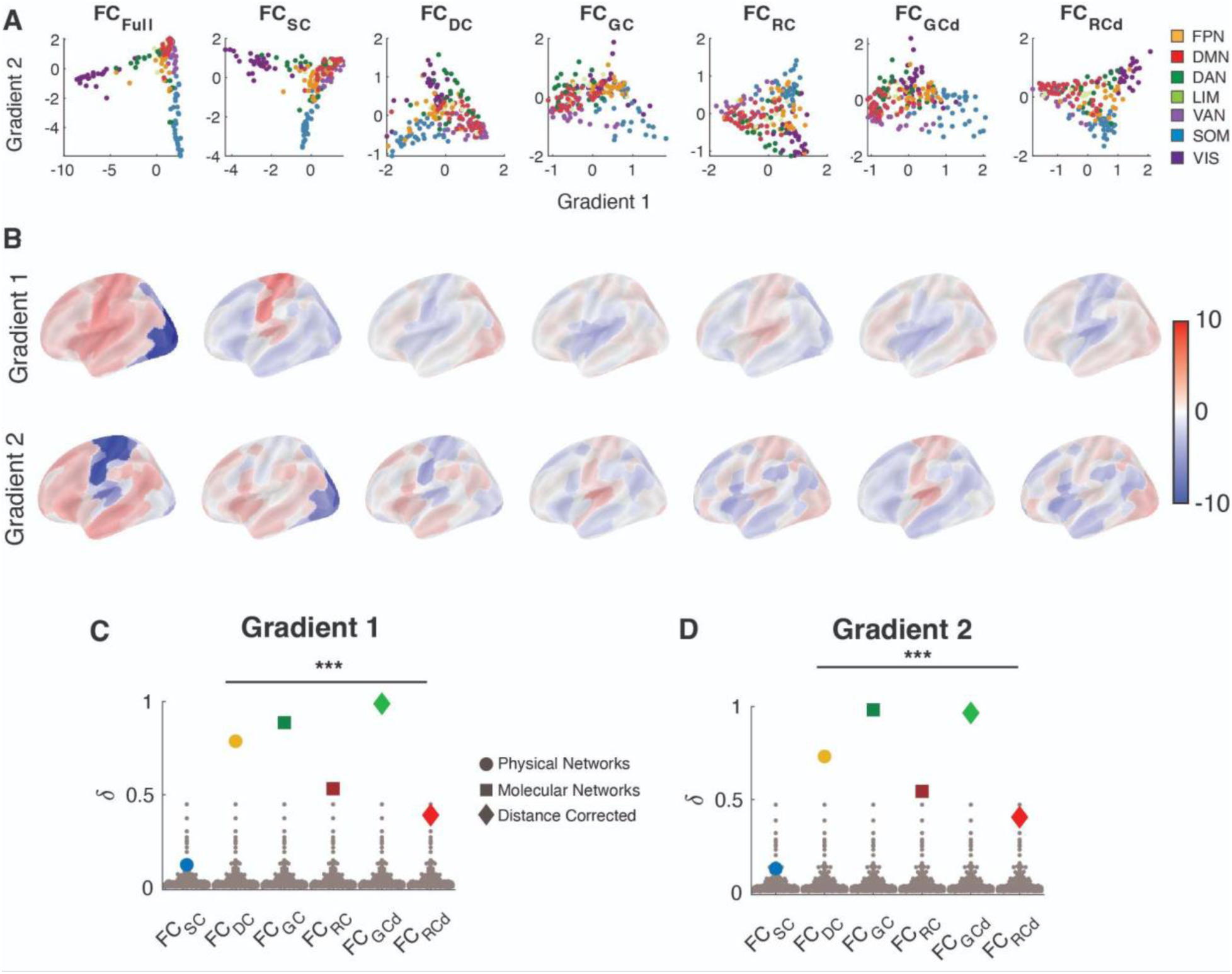
Biophysical networks shape the gradient organization of functional brain connectivity. (A) The first two connectivity gradients estimated from the fully connected functional brain network (FC_Full_), and RFNs, i.e., after removing direct connections associated with SC, DC, GC and RC, GC_d_ and RC_d_. (B) Brain plots of Gradient 1 (*top*) and Gradient 2 (*bottom*). Shift in (C) Gradient 1 and (D) Gradient 2 for different RFNs. Given the complexity of the gradient metric, here to estimate the shift, we calculate the Pearson correlation (R) between the gradient map of FC_Full_ and RFNs and then calculate the shift δ as 1-|R| implying dissimilarity in gradient profiles. The gray dots represent the estimated change – dissimilarity – for the null model which was constructed by removing a set of random connections from the functional connectivity network FC_Full_. FPN, Frontoparietal Network; DMN, Default Mode Network; DAN, Dorsal Attention Network; LIM, Limbic; VAN, Ventral Attention Network; SOM, Somatomotor; VIS, Visual Network. Statistical testing assessed the deviation from the null model with *** denoting p-value < 0.001.

Removing GC and GC_d_ connections destroys the relative placement of different communities compared to FC_Full_ whereas the impact of removing SC is relatively low. Removing all other biophysical networks preserves some aspects of the organization (**Fig. 7A,B**). To quantify these observations, we estimated the shift between FC_Full_ and RFN gradients, δ, by first calculating the Pearson correlation (R) between the gradient map and then calculating δ as 1-|R|, implying dissimilarity in gradient profiles. We used |R| instead of R to account for the fact that gradients might be aligned in different directions leading to positive and negative values, whereas we are interested in the overall extent of the deviation. We also compared the δ estimated from each RFN to a null model in the same manner as was done on the graph-theoretic measures.

Reflecting the qualitative observations, GC had the most pronounced effects on both Gradients 1 (δ = 0.88, P_rand_ < 0.001; **Fig. 7C**) and Gradient 2 (δ = 0.98, P_rand_ < 0.001; **Fig. 7D**). In fact, correcting for distance dependencies in GC increased the effect. DC also showed a strong effect on both Gradients 1 (δ = 0.78, P_rand_ < 0.001; **Fig. 7C**) and Gradient 2 (δ = 0.73, P_rand_ < 0.001; **Fig. 7D**). On the other hand, the effect of RC networks was substantially lower while the effect of SC is below significance (P_rand_ > 0.05) level. These observations imply that distance and genetics play a pivotal role in shaping the observed gradient organization in FC.

To determine the robustness of these results, we tested if the observed differences in RFNs was a function of the percent variance explained by each gradient between different RFNs. As can be observed in **Fig. S2A**, we did not observe any substantial differences in the percent variance explained by each gradient between RFNs, indicating that the observed shifts were not driven by changes in the amount of variance explained between gradients, but can in fact be attributed to different underlying biophysical factors.

A strong dependence of gradients on GC indicates that gradient organization could reflect genetic relationships between brain regions. However, the prominence of DC suggests that confounds such as Euclidean distance may also drive overall gradient organization of the functional connectivity. Therefore, caution is warranted when attributing gradient organization solely to hierarchical cognitive architecture. Nonetheless, unlike graph-theoretical measures, which were completely dominated by molecular factors, we see a substantial influence of a physical factor when higher-order interactions are considered.

### Unique contribution of underlying biophysical networks to functional connections

The strong concordance between FC and biophysical networks raised the question of whether a functional connection was supported by multiple biophysical factors. To determine if multiple biophysical factors underly functional connection, we ranked the edges within each biophysical network so that stronger connections had a higher rank and then averaged the ranks across networks. The new network was then thresholded to match the density of the other biophysical networks and an RFN was created from this new combined biophysical network. We conducted this procedure with and without including SC connections in the ranking as the sparsity of the SC network could skew the ranks. The analysis revealed that the combined biophysical network (FC_BP_) and without SC associated connection (FC_BPsc_) exerted minimal effects on both graph-theoretic and gradient properties indicating that biophysical networks largely support different functional connections (**Fig. S3**).

### Comparable impact of removing biophysical and strongest functional edges on FC organization

Additionally, the strong concordance between FC and biophysical networks raised the question pertaining to how to FC connections that share an underlying biophysical substrate compare to connections that do not. Previous work has shown that FC edges that share an underlying structural connections are stronger than unconnected edges (Honey et al., 2009). As can be observed in **Fig. S4A**, this relationship extends to the other biophysical networks (P < 0.001). The strength of connections associated with molecular factors, especially RC, were stronger than all others and SC associated connections were the weakest.

However, it remains unknown how removing edges in FC that shared an underlying biophysical network compare to removing the strongest edges in FC. This comparison is an important benchmark for determining the proportion of FC organization shaped and constrained by each biophysical factor. Therefore, we compared the observed percent change in graph-theoretic and gradient properties associated with each biophysical factor to an FC network that had the top 16.07% of the strongest connections removed, corresponding to the density of each biophysical network in our main analysis.

The results indicated that the top biophysical factor – RC – accounts for at most half (Ratio_RC_ = 0.48) of the changes observed in graph-theoretic properties after removing an equivalent number of the strongest connections in the FC (**Fig. S4B, C**). The ratio is estimated by comparing the changes induced by a biophysical network to the changes induced by removing the strongest functional connections. Extending the analysis to gradient organization, for Gradient 1, comparative effects were observed when removing GC associated connections in FC with removing an equivalent number of the top connections in FC (Ratio_GC_ = 1.01) and all other biophysical factors had weaker effects (**Fig. S4D**). Whereas, for Gradient 2, removing all other biophysical factors, except SC, had a stronger effect than removing the equivalent strongest connections in FC (**Fig. S4E**). The strongest effects were observed after removing GC associated connections (Ratio_GC_ = 2.48). These results indicate that the impact of biophysical networks is comparable to removing the strongest functional connections in FC.

### Robustness to methodological choices

Our main analysis on graph-theoretic and gradient organization utilized a single thresholding value based on the density of the SC network. To ensure that our results are robust across a range of thresholds, we repeated the analysis with thresholds that removed a range from 5 to 50% of connections in the FC. Reflecting our main results, for most thresholding values, molecular (GC and RC) based biophysical networks consistently had the strongest effects for the graph-theoretic properties of FC (**Fig. S5A)**. Whereas, DC had the strongest effects when more than 40% of connections were removed and SC consistently induced the weakest effects. Similarly for the gradient properties, DC and GC induced the strongest effects, but the extent of the effect varied with the percentage of connections removed (**Fig. S5B)**.

Further, our analysis was based on thresholding FC, but the FC network can be thresholded in any number of ways and differences in organization will be observed. To ensure that the differences we observed were meaningful and not simply due to thresholding, we compared the observed differences against three different types of thresholding, null models: (1) removing random connections (**Fig. 5 and Fig. 7**); (2) equating the number of intra- and inter-hemispheric connections removed (**Fig. S6**); and (3) degree preserving random connections (**Fig. S7**). Across the different null models, the effects induced by biophysical networks were similar and significantly greater (P < 0.001) indicating that the observed differences in FC organization due to biophysical factors were not simply due to removing connections at random.

Specifically, one large organizational difference between physical networks (SC, DC) and correlation matrices (GC, RC. FC) is that in correlation matrices, homotopic connections tend to be stronger whereas in physical networks, homotopic connections tend to be weak or absent (Hansen et al., 2023). Therefore, it is important to determine how much are graph theoretical measures of FC affected simply due to the removal of many homotopic connections? We tested for this by designing a null model that removes the same number of inter- and intra-hemispheric edges as were removed from the empirical network. We obtained similar results as our main findings in that molecular factors were significantly different even after accounting for homotopic differences for both graph-theoretic and gradient properties (**Fig. S6**).

Additionally, we examined the robustness using a different null-model where instead of removing connections from FC_Full_ at random. Specifically, the connections in each biophysical network were randomized while preserving the degree distribution (Maslov and Sneppen, 2002). As discussed in **Fig. S7**, we obtained similar results for the re-wired null model for both graph-theoretic and gradient properties replicating all our main findings discussed in Figures 4 and Figure 7.

Thus far, the analysis on graph-theoretic and gradient organization was based on the magnitude (i.e. |correlation|) of the FC. To test that this choice did not affect our results, we conducted an additional analysis where we first thresholded the FC at zero to remove all negative connections (Schwarz and McGonigle, 2011; Zhan et al., 2017). We observed differences in the magnitude of network features, as expected, however, we observed a clear dominance of the effect of molecular factors on graph-theoretical features replicating our main findings (**Fig. S8A,B**).

Similarly, we observed a prominent (i.e. strongest) effect of DC on gradients (**Fig. S8C,D**). The effect of molecular factors was somewhat nuanced; GC showed a dominant effect on Gradient 2, as discussed in Figure 7, but Gradient 1 had a higher impact of RC as compared to both GC and DC.

### Generalization of findings across datasets

Additionally, we corroborated our main findings using additional resting-state fMRI data recorded from 130 healthy individuals from the LA5c dataset (Poldrack et al., 2016). The goal of the LA5c dataset was to understand the dimensional structure of memory and cognitive control in both healthy individuals and individuals with neuropsychiatric disorders including schizophrenia, bipolar disorder, and attention deficit/hyperactivity disorder. For the corroborative analysis, we utilized approximately 5 minutes of resting-state fMRI data in the healthy population and the results are described in **Figure S9 and Figure S10**.

Similar to our findings with the HCP data (**Fig. 4**), biophysical networks exerted the strongest effects on the modularity (Δ = 39.67 ± 16.45%; **Fig. S9A**). Moreover, FC was most strongly constrained and shaped by molecular factors (GC: Δ = 20.52 ± 15.73% and RC: Δ = 19.41 ± 18.86%) reflecting the findings in the HCP dataset (**Fig. S9B-I**). While molecular factors dominated in both HCP and LA5c, we observed a relatively higher impact of genetic similarity between brain regions (GC) in LA5c than in the HCP dataset in which RC was the dominant molecular factor. However, as was found in the HCP dataset, SC had the least effect on FC organization. Further, the findings from the gradient analysis revealed a dominance of DC and GC on Gradient 2, and DC on Gradient 1 (**Fig. S10**), similar to the findings discussed in Figure 7. However, we found a more nuanced effect of molecular factors on Gradient 1. Additionally, we observed a significant effect of SC on gradients, unlike what was observed in HCP data.

Across these various corroborating analyses, we observed a clear dominance of molecular factors on the graph-theoretical organization of functional connectivity. Some of the gradient features showed variations across the datasets, but the dominant effect of DC was robustly observed on the gradient organization of functional connectivity for different methodological and data choices considered in this study.

### Predicting FC from underlying biophysical networks

The above analysis indicates that molecular factors constrain and shape the pairwise relationships between brain regions. Therefore, as an additional analysis to establish to corroborate our findings, we determined how well each of the biophysical networks could predict FC. We observed that the biophysical networks exhibited differential capabilities when predicting FC (**Fig. S11**). On average the best predictor of nodewise functional connectivity was neuroreceptor congruence (R^2^ = 0.31 ± 0.16; P < 0.001). Whereas the other biophysical networks exhibited weak or localized predictive capabilities. Moreover, neuroreceptor congruence was the best predictor of functional connectivity in the healthy control group from the LA5c dataset, reflecting the results in the HCP dataset (**Fig. S12**). Together these results confirm the dominance of molecular factors in constraining and shaping functional connectivity across the cortex.

### Biophysical networks underlie neuropsychiatric induced changes in FC

Lastly, we investigated if specific biophysical networks underlie the changes in functional connectivity observed in neuropsychiatric populations. The deviations in the case of neuropsychiatric conditions tend to be more nuanced (Segal et al., 2023; Winter et al., 2022). We aimed to uncover if these changes can be attributed to different biophysical factors to a varied extent. We leveraged the LA5c dataset which also included individuals with neuropsychiatric disorders – schizophrenia (N=50), attention deficit/hyperactivity disorder (N=40), and bipolar disorder (N=37) (Poldrack et al., 2016). The strength of the LA5c dataset is that it allowed us to include a diverse set of neuropsychiatric disorders while minimizing the issues related to experimental protocol and measurement differences.

First, we tested if the functional connectivity tends to increase or decrease in the clinical groups. We performed an edgewise independent sample t-test to identify connections that changed due to schizophrenia (SCZ), attention deficit/hyperactivity disorder (ADHD) and bipolar disorder (BD) compared to healthy individuals, respectively. We observed that some connections increased (red) and others decreased (blue) in strength compared to healthy individuals (**Fig. 8**) with overall functional connectivity increasing in SCZ (average t-value = 0.30 ± 0.009; **Fig. 8A**) and decreasing in ADHD (average t-value = -0.35 ± 0.007; **Fig. 8D**) and BD (average t-value = -0.47 ± 0.007; **Fig. 8G**). We then investigated whether these alterations were shaped significantly by specific biophysical networks.

**Figure 8.**
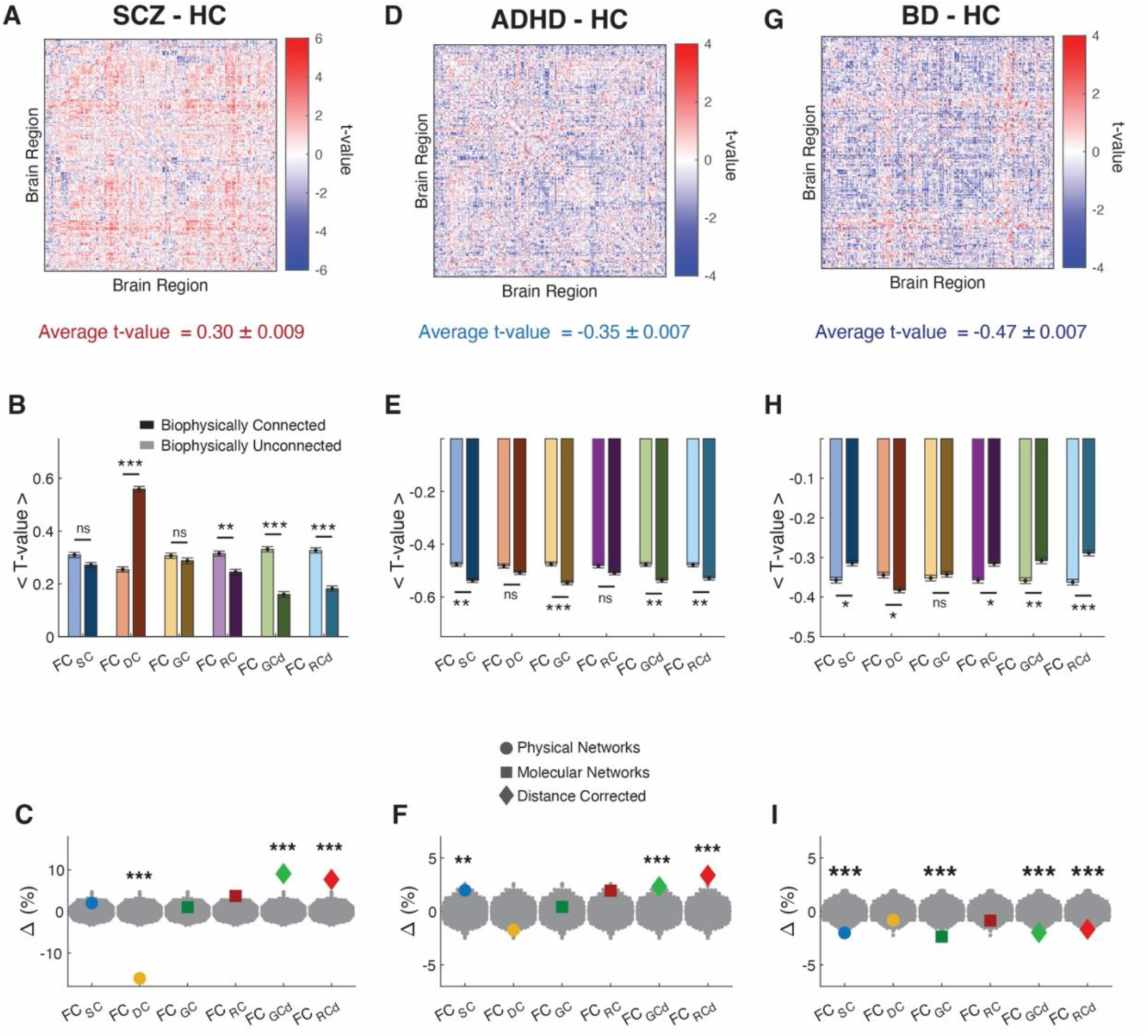
Neuropsychiatric disorders induced changes in functional connectivity are related to biophysical networks. (A) Differences in functional connectivity (independent samples t-tests) between schizophrenia (SCZ) and healthy controls (HC) with average t-value shown at the bottom. (B) Average t-value of FC connections that are connected or unconnected in an RFN. (C) Biophysical networks induced percent change in average t-value. We constructed RFNs from the t-value matrix by removing connections specific to each biophysical network and computed the shift Δ as a percent change in average t-value. (D-F) Same as panel A through C, but for attention deficit/hyperactivity disorder (ADHD). (G-I) Same as panel A through C, but for bipolar disorder (BD). Statistical testing assessed observed differences in RFNs compared to a null model (gray dots) based on removing an equal number of random connections. *** P < 0.001; ** P < 0.01; * P < 0.05; ns, not significant.

First, we compared all connections not in the RFN to those in the RFN because this would provide an estimate of the relationship between observed changes in psychiatric illness with underlying biophysical factors. We estimated the average t-value for connections that shared an underlying biophysical connection with those that did not.

As can be observed in **Figure 8B**, for schizophrenia, connections that shared a DC associated connections exhibited a larger increase than unconnected, but the trend was reversed for all other biophysical networks (P < 0.01). For bipolar disorder, in general, connections that shared an underlying biophysical connection exhibited larger decreases in strength (P < 0.01; **Fig. 8E**). Similar trends were observed for ADHD with connections broadly decreasing in strength (**Fig. 8H**).

To confirm these findings, we calculated RFNs from the t-value matrix by removing connections associated with each biophysical network in the same manner as described in Figure 1 and quantified the deviation (Δ) as a percent change in average t-value. A positive Δ indicates that connections associated with a specific biophysical network acted to decrease average FC (i.e., negative t-values) in the clinical population as compared to healthy controls and upon removal, the average t-value increased in the corresponding RFN. Similarly, a negative Δ indicates that connections associated with a specific biophysical network acted to increase average FC in the clinical population. Moreover, we compared the observed changes to a null distribution based on removing an equal number of random connections in the same manner as was conducted for the above analyses.

For schizophrenia, removing Euclidean distance dependent connections (DC) resulted in significant decrease in Δ (i.e., -16.10%, P_rand_ < 0.001) indicating that DC associated connections were heightened in strength in the schizophrenia group compared to healthy controls (**Fig. 8C**). We did not observe a significant effect of removing SC and molecular factors, i.e., GC and RC (P_rand_ > 0.05), whereas, removing the molecular connections corrected for distance, i.e., GC_d_ (Δ = 9.07%), RC_d_ (Δ = 7.68%), resulted in significant increase in Δ (P_rand_ < 0.001) indicating that these networks dampened functional connectivity in SCZ.

In ADHD, we observed a similar positive effect of distance corrected molecular networks, i.e., GC_d_ (Δ = 2.32%; P_rand_ < 0.001) and RC_d_ (Δ = 3.38%; P_rand_ < 0.001), indicating that these networks dampened functional connectivity, however we found this effect to be much smaller than in case of SCZ (lower Δ values; **Fig. 8F**). Similar to SCZ, in ADHD too we did not observe a significant effect of GC and RC, however, unlike in SCZ, SC showed a low but significant positive effect (Δ = 1.97%; P_rand_ < 0.001), indicating that SC related connections are dampened in functional connectivity in the ADHD group. We did not observe a significant effect of DC for the ADHD group.

In the BD group we also observed a significant but opposite effect of SC (Δ = -2.00%; P_rand_ < 0.001), indicating that SC related functional connections were higher in this clinical group as compared to healthy controls (**Fig. 8I**). We also observed a significant negative effect of genetic similarity in both GC (Δ = -2.35%; P < 0.001) and GC_d_ (Δ = -1.97%; P_rand_ < 0.001) indicating that connections associated with genetic similarity were heightened in the BD group as well. Additionally, RC_d_ also led to significant negative change (Δ = -1.66%; P_rand_ < 0.001) while DC associated connections did not have a significant effect (P_rand_ > 0.05).

Our framework reveals interesting dependencies of the observed alterations in functional connectivity on underlying biophysical networks. We observed that SCZ and ADHD exhibited a similar relationship between molecular and distance corrected factors and functional connectivity. However, their relationship with physical factors, both SC and DC, differentiated these groups; SCZ is characterized by a strong negative shift due to DC and ADHD is characterized by a positive shift due to SC. On the other hand, BD exhibited a different profile of shifts from SCZ and ADHD across all biophysical networks, with SC and GC driven negative shifts uniquely separating it from the other groups. Overall, we found the effect of molecular factors to be more nuanced across all the groups and a strong dominance of physical factors in differentiating groups.

## Discussion

Functional brain connectivity is the cornerstone of cognition, with alterations potentially serving as indicators of pathological states (Fornito et al., 2015). This large-scale property of the brain, which also provides a window into brain-wide communication, is anchored by various underlying biophysical factors. Here we addressed how various biological factors, physical and molecular, shape the fundamental organizational features of the brain’s functional connectivity (FC). Specifically, we considered structural connectivity (Honey et al., 2009), Euclidean distance (Shinn et al., 2023), gene expression similarity (Richiardi et al., 2015), and neuroreceptor congruence (Hansen et al., 2022a) across regions and dissociated their individual contribution to pairwise (graph-theoretic) as well as higher-order (gradients) emergent properties of functional connectivity. We developed a computational framework leveraging remnant functional networks to determine the contribution of biophysical networks in shaping functional connectivity and further assessed how these factors drive the functional connectivity alterations observed in groups with neuropsychiatric disorders.

A critical question in neuroscience is understanding how the intricate brain wide communication is shaped and constrained by multiple underlying factors from physical to neurochemical factors. However, most brain network models assume that brain regions are homogenous and discard the neurobiological heterogeneity (Bazinet et al., 2023). Additionally, previous work has shown that both brain structure and function strongly depends on the underlying chemoarchitecture (Hansen et al., 2022a). In fact, the molecular features, both similarity in gene expression and neuroreceptor congruence, map onto brain network features such as rich club organization (Hansen et al., 2023). Moreover, similarity in gene expression and neuroreceptor congruence, can explain the cortical abnormalities in multiple neurological and psychiatric illnesses (Hansen et al., 2022a, 2022b).

In order to fully unravel the role of biophysical networks, we leveraged a comprehensive set of quantitative features derived from graph theory and topology, two branches of mathematics that have been instrumental in understanding the foundational features in complex neuroimaging data (Centeno et al., 2022). With the help of these features, we quantified and compared the organization of functional connectivity networks and remnant functional networks that were constructed from functional connectivity after removing connections associated with a given biophysical network. Our findings suggest that molecular factors, i.e., neuroreceptor congruence and similarity in gene expression between regions may play a stronger role in shaping the graph-theoretic organization of the resting-state functional brain connectivity estimated with fMRI than physical factors. Moreover, our results suggest that gradient properties are dependent on both physical (i.e. Euclidian distance) and molecular (i.e. genetic similarity) factors.

In particular, neuroreceptor congruence among brain regions shaped a wide spectrum of graph-theoretical properties that have been discussed to capture a variety of functions in the brain including efficient processing of information (Avena-Koenigsberger et al., 2017), robustness to malfunction (Aerts et al., 2016) and global excitation (Bansal et al., 2018), and even the emergence and termination of seizures (Rungratsameetaweemana et al., 2022). In general, we observed a higher shift in graph-theoretical features, such as weighted degree and clustering coefficient, after removing molecular factors, and we found a unique relationship between the magnitude of the shift with both weighted degree and clustering coefficient for neuroreceptor congruence (Fig. 6 and Fig. S1) implying it may play a role in shaping hub-like regions in the brain which underlie efficient integration and segregation of information (Cohen and D’Esposito, 2016). Additionally, using search information-based modeling (Fig. S12 and Fig. S13) we found that the neuroreceptor congruence was the best predictor of functional connectivity (Goni et al., 2014). This reliance on neuroreceptor congruence could suggest that pairwise interactions in functional brain connectivity emerge from a similar response to neurotransmitters across the brain.

Differing from graph-theoretic features, we found a mix of physical and molecular factors, Euclidean distance and similarity in gene expression, to shape the gradient features of functional connectivity capturing higher-order interactions. The shared genetic profiles may provide a foundation for the gradient properties that are linked to the functional specialization or hierarchy observed in the brain networks (Margulies et al., 2016). However, the dependence on Euclidean distance we observed across different methodological choices and datasets may suggest a more physical basis of gradient organization. These results also reflect previous findings of Watson and Andrews, 2023 showing that gradients identified by connectopic mapping techniques reflect confounds associated with Euclidean distance. Irrespective of the overall interpretation of the gradient organization, our analysis suggests that Euclidean distance may drive the higher-order interactions in the functional brain connectivity.

In line with previous work, our results suggest that a linear mapping of structural connections provides a limited understanding of the structural underpinnings of the brain’s functional connectivity (Fotiadis et al., 2024). Structural pathways are undoubtedly critical for facilitating communication between neurons and maintaining healthy brain states (Warren et al., 2014), but our study highlights the limitations when inferring functional connectivity directly from structural connections estimated with diffusion MRI (Sotiropoulos and Zalesky, 2019). Factors such as the composition of the neuroreceptors on dendrites can amplify signals from weak structural connections or diminish signals from strong connections, thus making it difficult to infer the extent of functional connectivity from structural connections alone (Harnett et al., 2012). This weak relationship could be because SC does not linearly map onto FC and nonlinear mapping can improve structure-function relationships (Honey et al., 2010, 2009). Moreover, it is important to note that the analysis on SC was based on group-averaged SC. However, averaging SC connections across individuals may bias the group-averaged SC towards short-range connections. As a result, this may dampen the observed influence of SC on the organization of FC. This bias toward short-range connections may be overcome using distance-dependent consensus thresholding to account for the long-range connections (Betzel et al., 2019). Taken together, these findings suggest that for a more holistic understanding of how functional connectivity arises, we need to refine the modeling frameworks that largely rely on structural connectivity, and we need to better understand how genetic expression and neuroreceptor congruence across the cortex shapes brain-wide communication.

One can generate many different types of brain networks from multimodal imaging. Besides functional, physical and molecular networks examined here, as previously shown (Hansen et al., 2023) brain networks can be generated from laminar organization and metabolic similarity. Our findings complement the observations in in which they observed that molecular factors shape and constrain the organization of brain network features (Hansen et al., 2023). Additionally, our analysis provides a framework to compare across different networks.

Further, using our framework we determined the factors that may contribute to altering functional connectivity in neuropsychiatric disorders. Consistently in schizophrenia, bipolar disorder, and ADHD, complex interaction between biophysical networks underlined the observed changes in functional connectivity. However, we observed that physical factors differentiated disorders. In particular, we observed a strong impact of Euclidean distance uniquely in schizophrenia suggesting that connections associated with distance dependent connectivity were heightened. Whereas, structural connectivity driven differences separated all the groups, such that SC related functional connections were enhanced in bipolar disorder, unaffected in SCZ, and dampened in ADHD as compared to healthy controls. Further studies are necessary to validate these findings in bigger datasets as well as to assess regional specificity in connectivity alterations. Nonetheless, our analysis provides specific targets and hypotheses to develop and test disease specific diagnosis and perhaps treatment.

Some considerations are warranted while interpreting our findings and designing follow up analysis. Importantly, the focus of the analysis has been on understanding the relationship between biophysical networks and functional connectivity at the macroscale, but it remains unknown whether the relationships observed here are reflected at a more microscopic scale of neurons and neural circuits. Moreover, for the analysis on GC, gene expression samples were mirrored across hemispheres, and this may result in artificially large homotopic connections. This limitation could be mitigated in future studies as more detailed gene maps become available. Additionally, our analysis focused on four distinct types of biophysical networks, but biophysical networks can be estimated from a multitude of other factors such as similarity in neuronal composition (Siletti et al., 2023) or cytoarchitecture (Paquola et al., 2020) between brain regions which might also impact the organization of functional connectivity. Nonetheless, our framework provides a means to assess the relationship between functional connectivity and multiple biophysical networks ranging from the micro (i.e., neurons) to the macroscale.

In conclusion, our study highlights that various biophysical factors, both physical and molecular, play a role in shaping the fundamental emergent properties of the brain’s functional connectivity. These factors should be accounted for while creating working models of functional connectivity and sole reliance on the structural connectivity should be avoided. However, the contributions of different biophysical factors may not be equitable, and these factors differentially shape distinct aspects of functional connectivity. Molecular factors, particularly neuroreceptor congruence, shape graph-theoretic properties that capture pairwise interactions in functional connectivity, while a mix of molecular and physical factors, particularly, Euclidean distance and similarity in gene expression drive the gradient properties of functional connectivity capturing higher-order interactions. Additionally, physical factors, including Euclidean distance and structural connectivity seem to better differentiate disease specific alterations. These findings can be factored in to understand and model functional connectivity in various specific applications.

While our present study focuses on the resting-state functional connectivity, our analysis provides a simple yet powerful test to further examine how various underlying factors uniquely and/or dynamically shape functional connectivity in varied domains including during task performance, development, and clinical.

## Methods

### Analytical framework

#### Remnant functional networks (RFNs)

We propose remnant functional networks (RFNs) to quantify the contribution of underlying biophysical networks in shaping functional brain connectivity. The critical idea behind the framework is that it quantifies and compares the change in the organization of functional networks when shared connections between the functional brain network and a biophysical network of interest, i.e., structural connectivity, Euclidean distance, similarity in gene expression, or neuroreceptor congruence, are removed (**Fig. 1**).

Specifically, first we estimate the functional connectivity between brain regions using the BOLD signal from functional MRI. This results in a fully connected weighted functional network (FC_Full_; **Fig. 1A**). Second, connections in FC_Full_ that also have a corresponding connection in the reference biophysical network (e.g., SC, DC, GC or RC) are removed (**Fig. 1B)**. For instance, when the reference network is SC, procedurally this entails setting all connections in the FC_Full_ network to zero that also have structural connections; this produces a weighted functional network with connections remnant to (indirect) structural connections (FC_SC_; **Fig, 1C**; note that the subscript corresponds to the connections that are removed). The same procedure is repeated for DC, GC and RC based networks resulting in weighted networks with only remnant connections, i.e., FC_DC_, FC_GC_, FC_RC_, respectively. To equate the density across biophysical networks, we estimated the average density of the SC network across the 48 subjects (d_ave_ = 16.07%) since it is the least dense compared to the other networks. Then each of the biophysical networks were thresholded so that the most prominent (i.e. strongest) 16.07% of connections were retained. These connections were subsequently removed from the FC to create the RFN with a density of 83.93%. Additionally, for the DC matrix, the thresholding process retains the shortest connections.

#### Quantifying the shifts in RFN

Critically, to determine which features of the functional network SC, DC, GC and RC shape, we quantify the magnitude of the change in the feature after removing SC, DC, GC or RC from the functional network. Specifically, a feature (e.g., network modularity) of the network is estimated in FC_Full_ which we label as P_Full_. Then, the same feature is calculated for RFNs, i.e., FC_SC,_ FC_DC_, FC_GC_, FC_RC_ networks, and for simplicity we label these as P_SC_, P_DC_, P_GC_, and P_RC_. Further, to assess the statistical significance, we constructed a null model by removing an equal number of connections from the functional connectivity at random (details below). We iterated on this process 1000 times and generated a null distribution.

### Functional MRI Data

#### HCP Resting-State Functional Connectivity

We used pre-processed resting-state functional MRI (fMRI) data from 48 healthy human participants from the Human Connectome Project (HCP) (Van Essen et al., 2013). We downloaded 50 participants, but two were missing data. In brief, participants underwent four sessions of 15-min resting-state scanning sessions. fMRI volumes were recorded using a customized 3T Siemens Connectome Skyra scanner with an EPI sequence (TR = 0.72 s, TE = 33.1 ms, 72 slices, 2.0 mm isotropic, field of view (FOV) = 208 × 180 mm). In addition to the pre-processing steps part of the HCP pipeline, we additionally regressed out the global signal. The data was mapped on Schaefer atlas to derive 200 ROIs (brain regions) (Schaefer et al., 2018). The same atlas was used in all subsequent biophysical networks as well (see below).

#### UCLA Consortium for Neuropsychiatric Phenomics Dataset (LA5c)

The Consortium for Neuropsychiatric Phenomics includes imaging of healthy individuals (N=122), individuals diagnosed with schizophrenia (N=50), bipolar disorder (N=37), and attention deficit hyperactivity disorder (N=40) (Poldrack et al., 2016). In brief, resting-state functional MRI scan were obtained while participants kept their eyes open for 304 s in the scanner. Neuroimaging data were acquired on a 3T Siemens Trio scanner. T1-weighted high-resolution anatomical scans (MPRAGE) were collected with a slice thickness = 1mm, 176 slices, TR=1.9s, TE=2.26ms, matrix=256 x 256, FOV=250mm. Resting-state MRI data were collected with a T2*-weighted EPI sequence with slice thickness = 4mm, 34 slices, TR=2s, TE=30ms, flip angle=90°, matrix=64 × 64, FOV=192mm.

The analysis was based on minimally preprocessed data using FMRIPREP version 0.4.4 (http://fmriprep.readthedocs.io). T1 weighted volume was corrected for bias field using ANTs N4BiasFieldCorrection v2.1.0 skullstripped and coregistered ICBM 152 Nonlinear Asymmetrical template. Resting-state data were motion corrected using MCFLIRT v5.0.9. Functional data was skullstripped and coregistered to the corresponding T1 weighted volume using boundary-based registration. Motion correcting transformations, transformation to T1 weighted space and MNI template warp were applied in a single step using antsApplyTransformations. Framewise displacement and Dvars were calculated using Nipype implementation. In addition to those regressors, global signal and mean white matter signal was also calculated and removed.

### Functional connectivity networks

For each individual, to extract functional connectivity we computed the magnitude of correlation in BOLD activity from a pair of regions across 200 regions derived from the Schaefer Atlas (Schaefer et al., 2018). For the HCP analysis a group functional connectivity network was estimated by averaging the individual subject networks together. The replication analysis using the LA5c dataset, a similar group functional connectivity network was estimated by averaging the individual subject networks together. The main analyses on the HCP dataset were based on the magnitude of the functional connections. We additionally replicated the findings in FC networks thresholded at zero in the HCP. Moverover, replication on the LA5c dataset was based on the magnitude of the functional connections.

### Biophysical networks

#### Diffusion MRI based structural connectivity (SC)

Preprocessed diffusion weighted images for each of the 48 subjects part of the HCP S1200 release were used to construct structural connectomes. Specifically, fiber tracking was done using DSI Studio with a modified FACT algorithm (Yeh et al., 2013). Data were reconstructed using generalized q-sampling imaging (GQI) in MNI space. Fiber tracking was performed until 250,000 streamlines were reconstructed with default parameters. For each subject, an undirected weighted structural connectivity matrix, A, was constructed from the connection strength based on the number of streamlines connecting two regions. Connectivity matrix was normalized by dividing the number of streamlines (T) between region *i* and *j*, by the combined volumes (*v*) of region *i* and *j*.

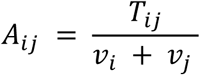

Group structural connectivity was estimated by averaging all the individual structural connectivity networks together. The group structural connectivity network was then thresholded to match the average density (d = 16.07%) across individual subjects.

#### Euclidean distance or distance derived connectivity (DC)

Functional connectivity between brain regions has been shown to be dependent on the physical proximity between regions, Euclidean distance. The distance between two regions was estimated as the Euclidean Distance (D) between their centroids. Since brain regions that are farther apart are going to have larger values, the matrix was converted to a proximity matrix (DC) by subtracting each entry from the largest value in the matrix as follow:

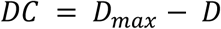

Where *D*_*max*_ represents the value associated with the farthest brain regions. For the DC matrix, the thresholding process retains the shortest connections.

#### Genetic expression based connectivity (GC)

Genetic expression dependent connectivity aims to estimate the extent of the similarity in genetic expression profiles between brain regions and relate the similarity in genetic expression to functional connectivity. Regional microarry expression data were obtained from 6 post-mortem brains (1 female, ages 24.0--57.0, 42.50 +/- 13.38) provided by the Allen Human Brain Atlas (AHBA, https://human.brain-map.org)(Hawrylycz et al., 2012). Data was processed with the abagen toolbox (version 0.1.3; https://github.com/rmarkello/abagen).

Microarray probes were reannotated using data provided by Arnatkevičiūtė et. al, 2019 (Arnatkevičiūtė et al., 2019); probes not matched to a valid Entrez ID were discarded. Next, probes were filtered based on their expression intensity relative to background noise (Quackenbush, 2002), such that probes with intensity less than the background in >=50.00% of samples across donors were discarded, yielding 31,569 probes . When multiple probes indexed the expression of the same gene, we selected and used the probe with the most consistent pattern of regional variation across donors (i.e., differential stability (Hawrylycz et al., 2015)), calculated with:

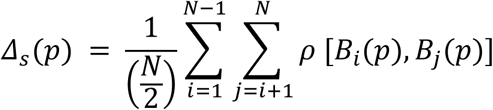

Where *ρ* is Spearman’s rank correlation of the expression of a single probe, *p*, across regions in two donors *B*_*i*_ and *B*_*j*_, and *N* is the total number of donors. Here, regions correspond to the structural designations provided in the ontology from the AHBA.

The MNI coordinates of tissue samples were updated to those generated via non-linear registration using the Advanced Normalization Tools (ANTs; https://github.com/chrisfilo/alleninf). To increase spatial coverage, tissue samples were mirrored bilaterally across the left and right hemispheres. Samples were assigned to brain regions in the provided atlas if their MNI coordinates were within 2 mm of a given parcel. All tissue samples not assigned to a brain region in the provided atlas were discarded.

Inter-subject variation was addressed by normalizing tissue sample expression values across genes using a robust sigmoid function (Fulcher et al., 2013):

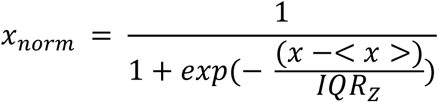

where < *x* > is the median and *IQR_z_* is the normalized interquartile range of the expression of a single tissue sample across genes. Normalized expression values were then rescaled to the unit interval:

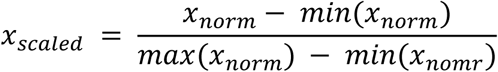

Gene expression values were then normalized across tissue samples using an identical procedure. Samples assigned to the same brain region were averaged separately for each donor and then across donors, yielding a regional expression matrix with 200 rows, corresponding to brain regions, and 15,633 columns, corresponding to the retained genes. Similarity in genetic expression (GC) between brain regions was estimated using the Pearson correlation.

#### Neuroreceptor congruence-based connectivity (RC)

Neuroreceptor dependent connectivity aims to estimate the extent of the similarity in neuroreceptor profiles between brain regions and relate the similarity in neuroreceptor congruence to functional connectivity. The analysis used data originally published in Hansen et al.,(Hansen et al., 2022). In brief, PET tracer imaging was used to map the density for 19 neuroreceptors and transporters. The neuroreceptors include 9 neurotransmitter systems: dopamine (D_1_, D_2_, DAT), norepinephrine (NET), serotonin (5-HT_1A_, 5-HT_1B_, 5-HT_2A_, 5-HT_4_, 5- HT_6_, 5-HTT), acetylcholine (*α*4*β*2, M1, VAChT), glutamate (mGluR_5_, NMDA), Gamma-aminobutyric acid (GABA_A_), histamine (H3), cannabinoid (CB1), and μ-opioid neuroreceptor (MOR). Each PET tracer image was parcellated onto 200 cortical regions and z-scored. A neuroreceptor similarity (RC) matrix was constructed using the Pearson correlation between the neuroreceptor profiles among the 200 cortical regions.

### Distance Correction

Since the similarity in gene expression and neuroreceptor composition between two regions can be dependent on the distance, we created two additional networks that controlled for distance.

We regressed the similarity due to Euclidean distance from the GC and RC networks to generate a distance controlled genetic network – GC_ED_ – and neuroreceptor network – RC_ED_.

### Density matching

The DC, GC, GC_ED_, RC and RC_ED_ are fully connected networks compared to the SC network which is sparsely connected, therefore, density matching was required to be able to compare the effect of removing each of these networks. We thresholded these networks to match the density of the SC network. As a result, the strongest connections survived in the biophysical networks.

### Graph-theoretical properties

We used multiple network properties to capture distinct aspects of the organization of functional brain connectivity. All properties were estimated using the Brain Connectivity Toolbox(Rubinov and Sporns, 2010). The specific properties measured are:

#### Weighted Degree

The weighted degree (*k_i_*) is defined as the sum of all connections for a brain region:

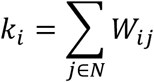

where *W* is the weighted adjacency matrix of a network with *N* number of brain regions(Rubinov and Sporns, 2010). The degree of nodes is an indicator of how important a node is for communication between brain regions(Rubinov and Sporns, 2010).

#### Clustering Coefficient

The weighted clustering coefficient (*CC*) for a brain region *i* is the intensity of triangles in a network calculated as,

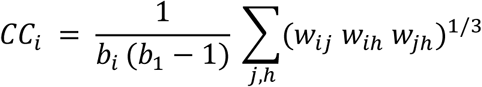

where *w* is the weighted adjacency matrix and *b* is the number of connections for brain region *i* and assess the extent of segregation in a network(Rubinov and Sporns, 2010).

#### Eigenvector Centrality

Eigenvector centrality (*EC_i_*) measures how influential a brain region is in a network, with a high value indicating a brain region is connected to other highly influential regions(Bonacich, 2007; Lohmann et al., 2010). The eigenvector centrality of a brain region *i* is given by the *i*-th entry in the dominant eigenvector, which is the vector ***v****=*[*v_1_,…v_N_*] that solves

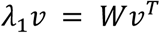

where *λ*_1_ is the largest eigenvalue of the weighted adjacency matrix, *W*. *Modularity*

Modularity (*Q*) quantifies the extent to which the brain functional connectivity may be subdivided into modules(Rubinov and Sporns, 2010). Specifically, for a given network *W*, *Q* is estimated as:

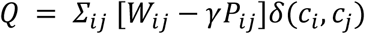

where *P_ij_* is the expected expected partition from a null model; *γ* is the resolution parameter (*γ =* 1); *c_i_* and c_j_ represent the module assigned to brain region *i* and *j* are; δ(c_i_,c_j_) is the Kronecker delta function and is equal to one when *c_i_* = *c_j_*, and is zero otherwise. Modules are thought to be a key element for cognition in which discrete processes occur(Bertolero et al., 2015).

#### Path Length

The characteristic path length (*L*) is the average shortest path length between all brain regions(Rubinov and Sporns, 2010),

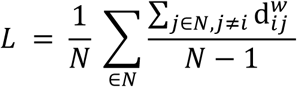

where is the distance between brain region *i* and *j*. To calculate, we first take the inverse of the edge weights to transform the weight to a measure of length (i.e., to transform a strong connection strength to a short length). The brain organization has been shown to be shaped by an economic trade-off between minimizing costs and allowing distant regions to communicate efficiently in order to adapt to changing environmental demands (Bullmore and Sporns, 2012).

#### Spectral Radius

The spectral radius measures the ease with which diffusion process can occur in a network. The spectral radius is calculated as,

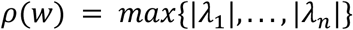

where |*λ*| corresponds to the absolute value of the eigenvalues of a network. Spectral radius reflects the excitability of brain networks(Bansal et al., 2018; Rungratsameetaweemana et al., 2022).

#### Synchronizability

Synchronizability is a measure of linear stability for a network of coupled dynamical systems(Motter et al., 2005),

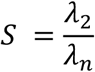

where λ_2_ is the second smallest eigenvalue of the unnormalized Laplacian matrix (L) and λ_n_ is its largest eigenvalue. The Laplacian is calculated as,

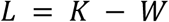

where K is the degree matrix of the functional connectivity matrix, W. Synchronizability assesses the ability of distant brain regions to synchronize(Arenas et al., 2008).

### Gradient organization

The hierarchical organization of functional brain connectivity is estimated using the framework based on gradients from Margulies *et. al.*, 2016 (Margulies et al., 2016). Gradients were estimated with the BrainSpace Toolbox (Vos de Wael et al., 2020). Specifically, FC_Full_ and RFNs were thresholded so that top 10% of connections were maintained and cosine similarity was used to estimate the similarity in the connectivity profiles between brain regions within each network, respectively. Analysis was based on the first two gradients identified using diffusion maps. All RFNs gradients were aligned to the FC_Full_ gradients using procrustes alignment.

### Search information-based modeling

Search information (SI) was used to model functional connectivity between brain regions. SI measures the amount of information (in bits) required to traverse the shortest paths between a given pair of nodes (Goni et al., 2014). The resulting SI matrix can be considered a proxy for the extent of functional connectivity between brain regions. Specifically, an SI matrix was estimated from each of the biophysical networks and the SI profile of each brain region was compared to the empirical functional connectivity profile (Liu et al., 2023). Note that higher (lower) SI values imply lower (higher) functional connectivity. In order to equate the relationship, the SI values were reversed by

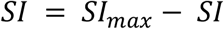

Where *SI*_*max*_ represents the value associated with the farthest brain regions.

### Biophysical networks underlying psychiatric illness

Edgewise independent sample t-test was used to identify connections that changed in the schizophrenia (SCZ), attention deficit/hyperactivity disorder (ADHD) and bipolar disorder (BD) compared to healthy individuals, respectively. The RFNs from the t-value matrix was estimated by removing connections associated with each biophysical network in the same manner as described in Figure 1 and the deviation (Δ) was quantified as a percent change in average t-value. The change was estimated between the average values in the fully connected T-value matrix and the average value in the RFN. Formally as,

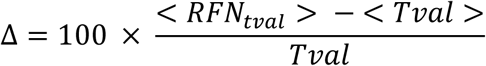

where < > corresponds to the average value in the *RFN*_*tval*_ and T-value matrix (*Tval*). Important to note that in this case the RFN is constructed from the T-value matrix.

### Null Models

Null model was used to benchmark the effects of removing SC, DC, GC and RC associated connections in the FC. Specifically, if a specific network feature in the FC is dependent on an underlying network, then the change in that feature after removing SC, DC, GC or RC associated connections should result in a change that is larger than what would be found with removing a random set of connections. To determine this, we used two different null models – random connections and re-rewired.

#### Random Connections

RFN null model was estimated by removing a random set of connections from the FC network that matched the number of connections in the biophysical network and the change in the network feature was calculated. This process was repeated 1,000 times to generate a null distribution for that feature.

#### Re-wired Null Model

To test the robustness of our results we used another null model that was estimated by removing a re-wired set of connections that preserved the degree of each node in the biophysical networks. This process was repeated 1,000 times to generate a null distribution for that feature. The re-rewired null model was generated using the Brain Connectivity Toolbox (Rubinov and Sporns, 2010).

## Data Availability

Minimally pre-processed Human Connectome Project data can be downloaded from https://www.humanconnectome.org/. Genetic expression data can be obtained using the abagen toolbox (https://abagen.readthedocs.io/en/stable/installation.html). Neuroreceptor composition data can be obtained at https://github.com/netneurolab/hansen_receptors. The brain connectivity toolbox can be downloaded at https://sites.google.com/site/bctnet/. Brainspace toolbox can be downloaded at https://brainspace.readthedocs.io/en/latest/pages/install.html.

## Acknowledgements

This research was supported by the U.S. Army DEVCOM Army Research Laboratory through mission funding (JOG) and army educational outreach program (KB, JN, W911SR-15-2-0001). The views and conclusions contained in this document are those of the authors and should not be interpreted as representing the official policies, either expressed or implied, of the U.S. Army DEVCOM Army Research Laboratory or the U.S. Government. The U.S. Government is authorized to reproduce and distribute reprints for Government purposes notwithstanding any copyright notation herein.

## Author Contributions

Conceptualization: JN, KB

Data Curation: JN

Formal analysis: JN

Funding acquisition: JOG, KB

Methodology: JN, KB

Supervision: KB, JOG

Visualization: JN, KB

Writing – original draft: JN, KB

Writing – review & editing: JN, JOG, KB

