## Supplemental Information for "Quantifying the influence of biophysical factors in shaping brain communication through remnant functional networks"

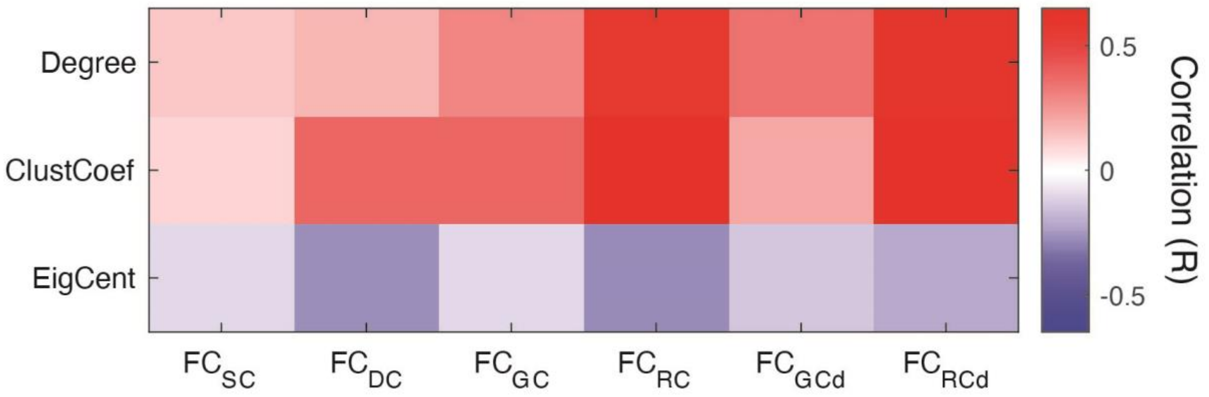


**Figure S1. Relationship between biophysical networks and regional network features.** The correlations between magnitude of change for each biophysical network across brain regions for weighted degree, clustering coefficient, and eigenvector centrality with strength of the feature. The correlation between $FC_{RC}$ and each of the network features is the same as in Figure 5D and is added for comparative purposes. Unlike RC driven networks, we did not observe a strong correlation between other biophysical networks and weighted degree.

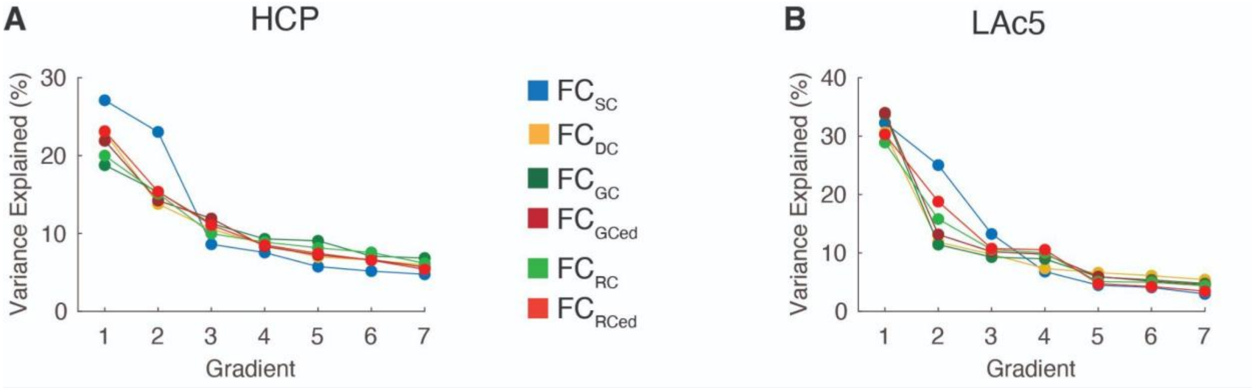


**Figure S2. Comparison of the variance explained by the first seven gradients for $FC_{Full}$ and for each of the RFNs**. (A) HCP and (B) LA5c datasets. Individual gradients explained comparable amounts of variance among RFNs.

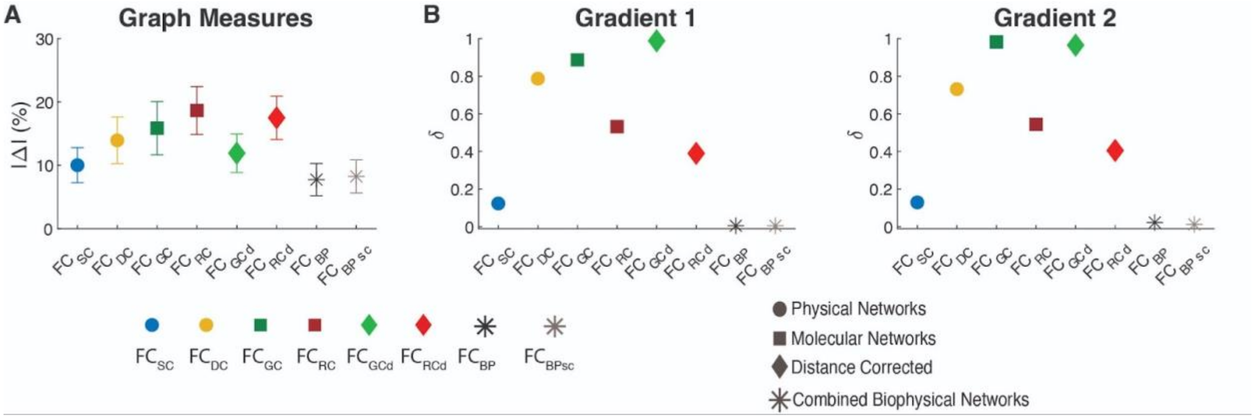


**Figure S3. Biophysical networks largely shape and constrain different aspects of FC.** Average change for biophysical networks across graph-theoretic metrics. For the analysis, the edges within each biophysical network were first ranked so that stronger connections had a higher rank and then averaged ranks across networks. The new network was then thresholded to match the density of the other biophysical networks and a RFN was created from this new combined biophysical network. The procedure was used to generate a combined biophysical network with and without ranked SC connections since the sparsity of the SC network might skew the ranks. The analysis revealed that the combined biophysical network (BP) and without SC associated connection ($BP_{sc}$) exerted minimal effects on both (A) graph-theoretic and (B) gradient properties. For comparative purposes we have included the changes from each of the biophysical networks from Figure 4B and Figure 7C,D.

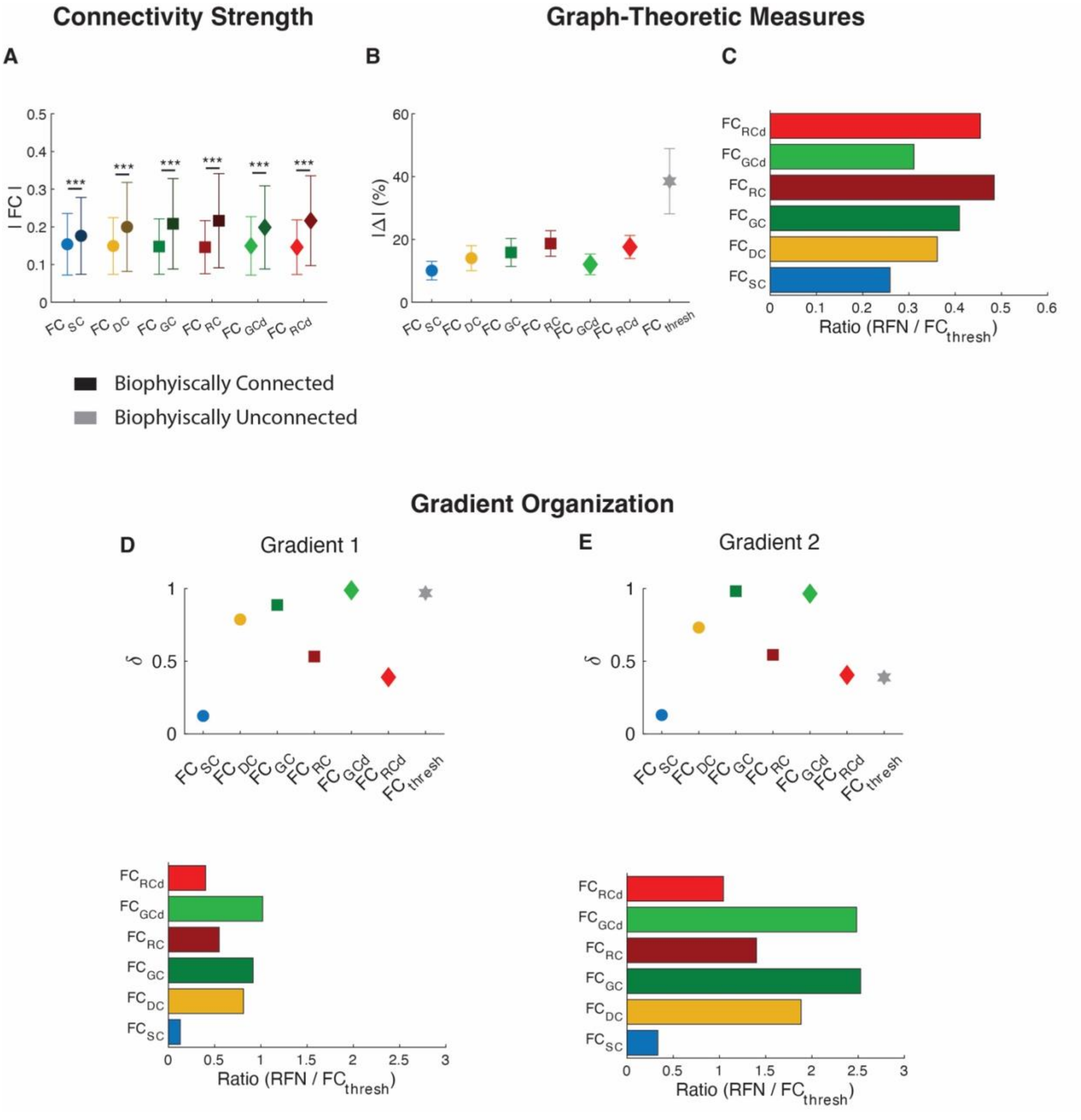


**Figure S4. Comparative impact of biophysical networks and strongest functional edges on FC organization.** (A) Strength of the functional connectivity between brain regions that are connected and unconnected by a biophysical network. First each biophysical network was threshold to a density of 16.07% and the connections in that remained were mapped onto FC. The analysis compared the magnitude of the FC strength between biophysically connected and unconnected brain regions. (B) Removing the top 16.07% of connections from FC ($FC_{thresh}$) induced the largest change in graph-theoretic properties compared to removing the equivalent percent of connections based on biophysical factors. Thresholding FC provides an upper limit of the expected percent change in graph-theoretic properties. The values for each biophysical factor are the same as in Figure 3B. (C) The ratio in average change in graph-theoretic between biophysical factors and thresholding FC. RC induced the greatest change with a ratio = 0.48 of the thresholded FC. (D) The change in Gradient 1 due to threshold FC is similar to the removing connections associated with genetic similarity (*top*). The ratio in average change in Gradient 1 between biophysical factors and thresholding FC (*bottom*). (E) Same as panel D, but for Gradient 2. The values for each biophysical factor are the same as in Figure 7C,D.

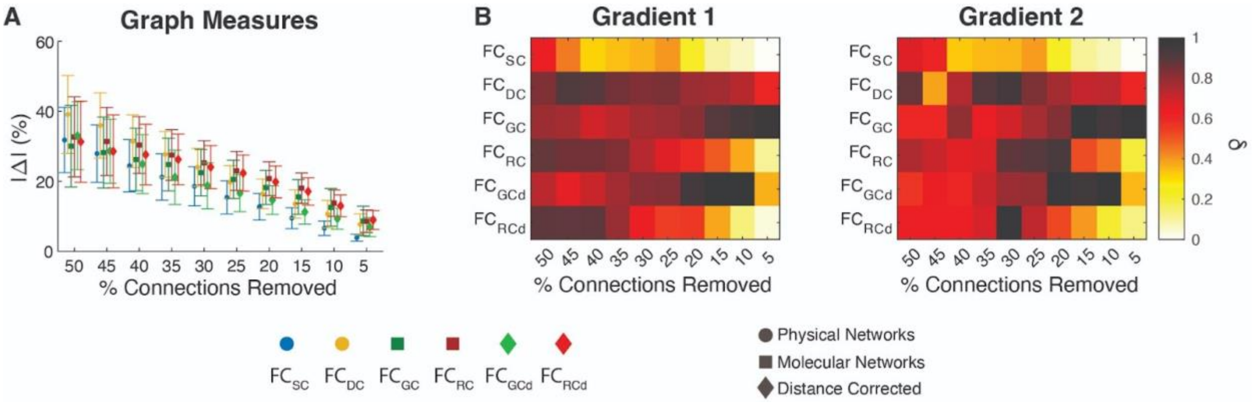


**Figure S5. Consistent relationship between biophysical networks, graph-theoretic and gradient properties across different thresholds in HCP dataset.** (A) Average change for biophysical networks across graph-theoretic metrics. Molecular (GC and RC) based biophysical networks consistently had the strongest effects for the graph-theoretic measures. Whereas, DC had the strongest effects when more than 40% of connections were removed and SC consistently induced the weakest effects. Note that results presented in the main analysis were based on removing the top 16.07% of connections in the biophysical networks. (B) For the gradient properties, DC and GC induced the strongest effects, but extent of the effect varied with the percentage of connections removed. To estimate the shift in the gradient properties, we calculate the Pearson correlation (R) between the gradient map of $FC_{Full}$ and RFNs and then calculate the shift δ as 1-|R| implying dissimilarity in gradient profiles. For both graph-theoretic and gradient properties, the thresholds ranged from 5 to 50% of connections removed.

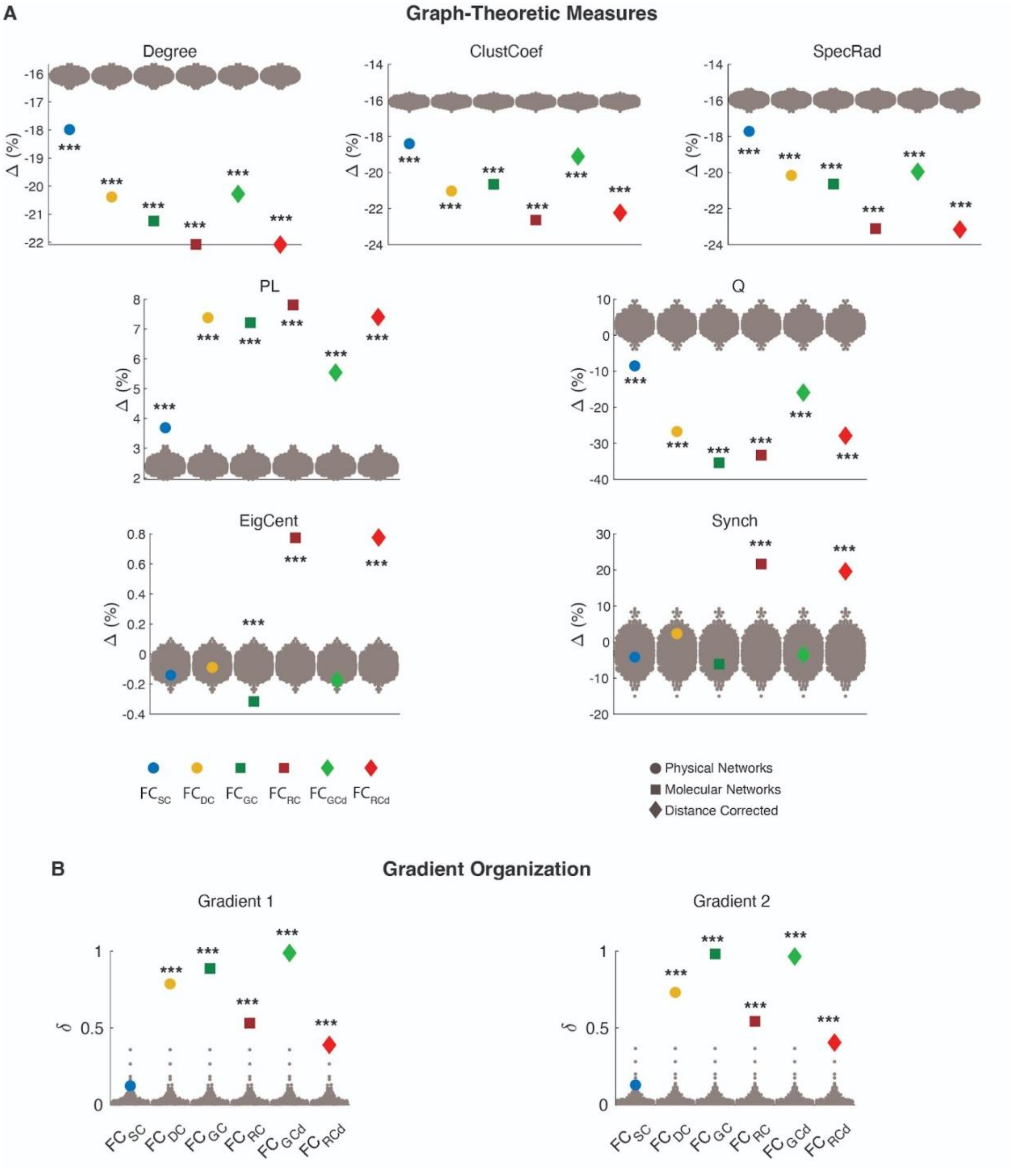


**Figure S6. Biophysical networks shaping the graph-theoretical and gradient properties of functional brain connectivity are not dependent on intra- and inter-hemispheric connections.** (A) Graph-theoretic and (B) gradient properties were significantly greater than a null model that removed the same number of inter- and intra-hemispheric (homotopic null) edges as were being removed from the empirical biophysical network. Gray dots represent the values after removing a homotopic null model random set of connections in the $FC_{Full}$ (null model). Statistical testing assessed the deviation from the null model with *** denoting p-value < 0.001.

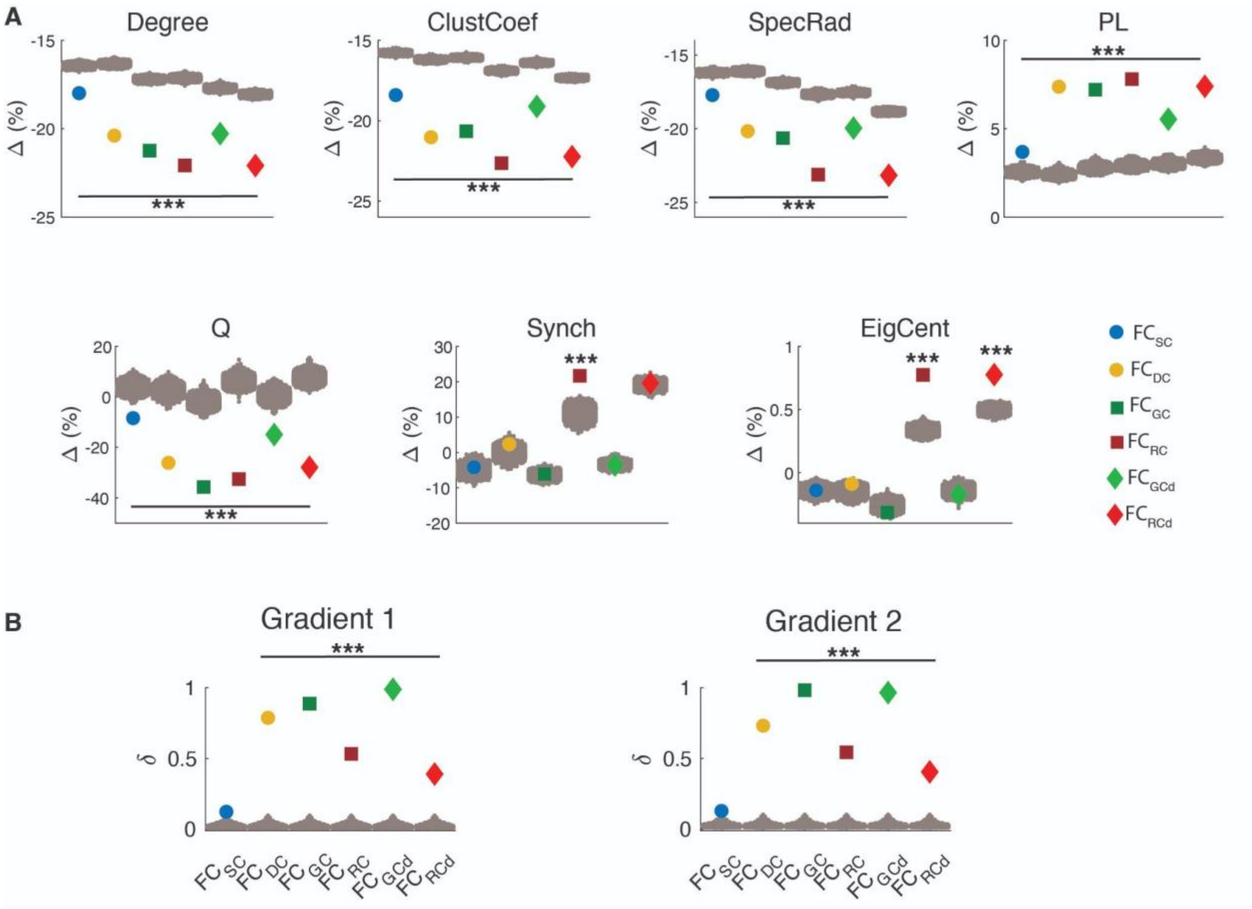


**Figure S7. Consistent relationships are obtained when using a null model based on re-wired biophysical networks.** (A) Graph-theoretic properties. Weighted Degree (Degree), Clustering Coefficient (ClustCoef), Spectral Radius (SpecRad), Path Length (PL), Modularity (Q), Synchronizability (Synch) and Eigenvector Centrality (EigCent). The effect of each biophysical network was quantified as a percent change (Δ) between $FC_{Full}$ and RFNs for each graph-theoretic measure. Colored dots represent the values after removing SC, DC, GC, RC, $GC_d$, and $RC_d$ associated connections in the $FC_{Full}$. (B) Gradient properties. Shift in Gradient 1 and Gradient 2 for different RFNs. Given the complexity of the gradient metric, here to estimate the change, we calculate the Pearson correlation (R) between the gradient map of $FC_{Full}$ and RFNs and then calculate δ as 1-|R| implying dissimilarity in gradient profiles.
Gray dots represent the values after removing connections in the $FC_{Full}$ from a re-wired, randomized biophysical network (re-wired null model). We generated 1000 total randomized networks for each biophysical network to construct the null distribution. Statistical testing assessed the deviation from the null model with *** denoting p-value < 0.001. Here, we observed a dominance of the molecular factors for the graph-theoretic measures (A) and a dominance of DC and GC associated connectivity for gradient features (B), which highlights the robustness of our findings discussed in Figures 4 and 6 for the choice of null model.

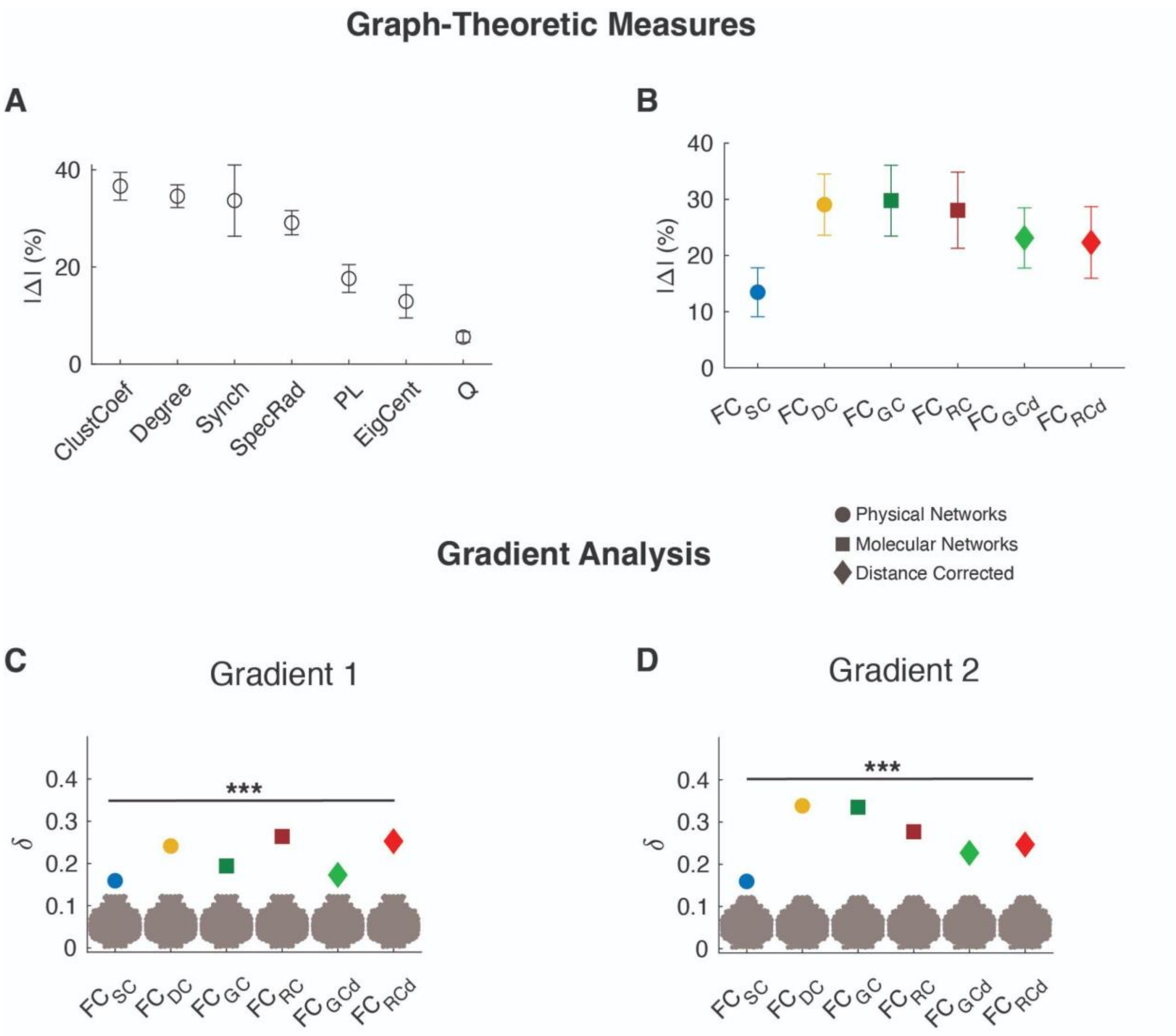


**Figure S8. The effect of biophysical networks is not dependent on the sign of connections in FC.** (A) Average change (Δ) in graph-theoretic metrics across biophysical networks. Functional connectivity estimated from the HCP dataset were thresholded at zero to remove negative connection prior to estimating graph-theoretic and gradient features. Due to the omission of negative edges, we observed changes in network properties when compared to Fig. 3, however, (B) average change for biophysical networks across graph-theoretic metrics shows the dominance of the molecular factors, as discussed in the main paper. (C) Shift in Gradient 1 and(D) shift in Gradient 2 for different RFNs. We calculate the Pearson correlation (R) between the gradient map of $FC_{Full}$ and RFNs and then calculate δ as 1-|R| implying dissimilarity in gradient profiles. The gray dots represent the estimated change – dissimilarity – for the null model which was constructed by removing a set of random connections from the functional connectivity network $FC_{Full}$. Statistical testing assessed the deviation from the null model with *** denoting p-value < 0.001. We find a significant effect of SC on both the gradients, and the effect of RC is also higher, particularly for Gradient 1, than discussed in Fig. 6. However, the effect of DC is one of the highest across all the biophysical networks, showing its robustness against this methodological choice.

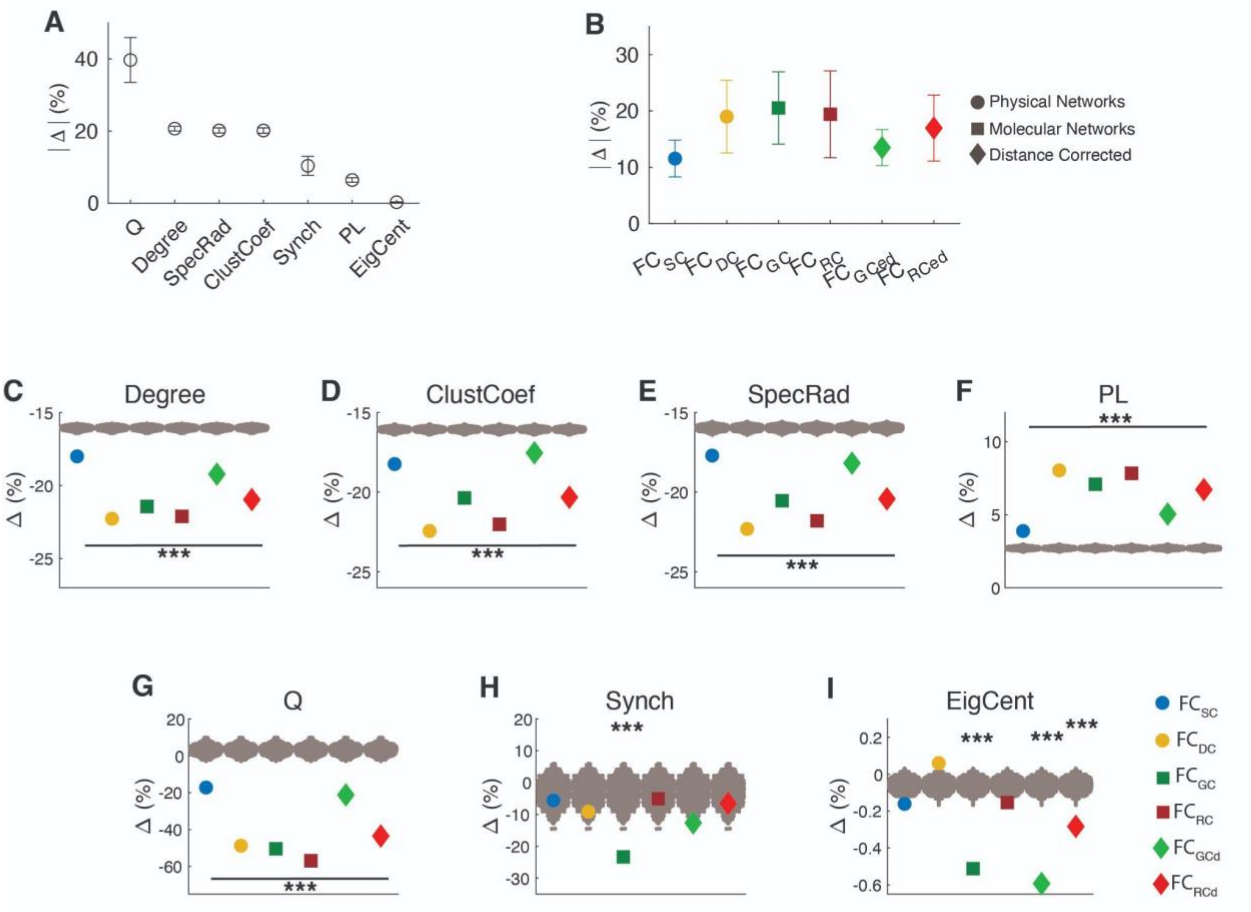


**Figure S9. Biophysical networks shaping the graph-theoretical properties of functional brain connectivity in the LA5c dataset.** (A) Average change in graph-theoretic metrics across biophysical networks. (B) Average change for biophysical networks across graph-theoretic metrics, highlighting the dominance of RC driven networks. (C) Weighted Degree (Degree), (D) Clustering Coefficient (ClustCoef), (E) Spectral Radius (SpecRad), (F) Path Length (PL), (G) Modularity (Q) , (H) Synchronizability (Synch) and (I) Eigenvector Centrality (EigCent).The effect of each biophysical network was quantified as a percent change (Δ) between $FC_{Full}$ and RFNs for each graph-theoretic measure. Colored dots represent the values after removing SC, DC, GC, $GC_d$, RC and $RC_d$ associated connections in the $FC_{Full}$. Gray dots represent the values after removing a random set of connections in the $FC_{Full}$ (null model). $FC_{SC}$: FC with SC associated connections removed; $FC_{DC}$: FC with DC connections removed; $FC_{GC}$: FC with GC connections removed; $FC_{RC}$: FC with RC connections removed; $FC_{GCd}$: FC with GC corrected for distance dependencies removed; $FC_{RCd}$: FC with RC corrected for distance dependencies removed. Statistical testing assessed the deviation from the null model with *** denotinges p-value < 0.001. Replicating the findings discussed in Figures 3 and 4, we observed a clear dominance of the molecular factors (GC and RC) on these graph-theoretic features while the effect of structural connectivity (SC) stays to minimum.

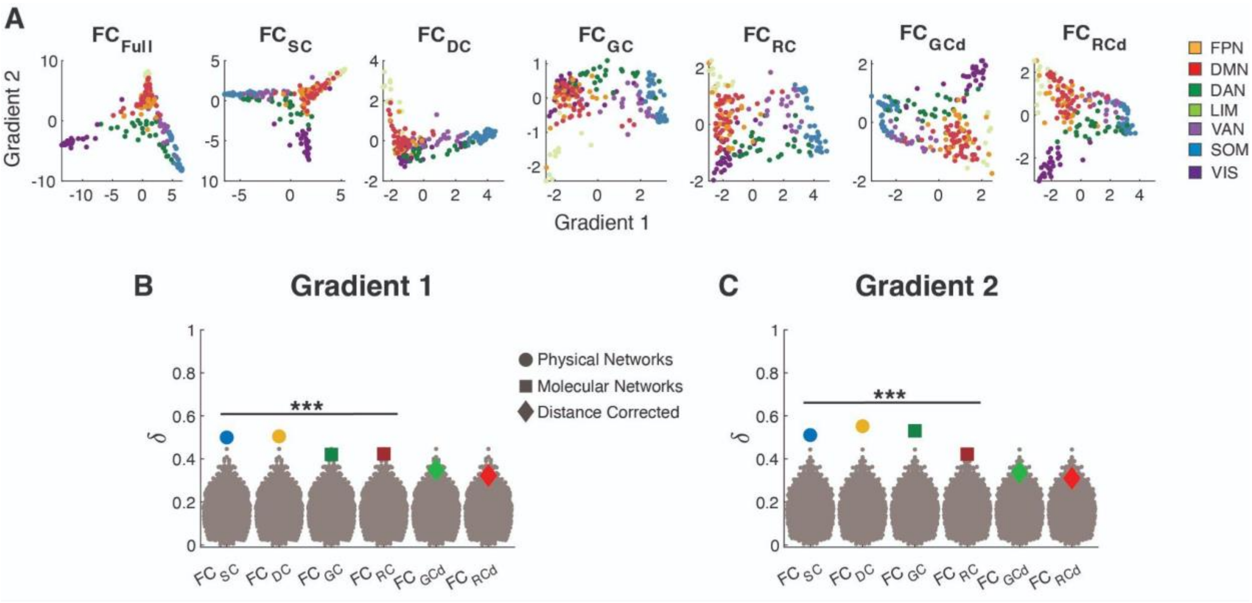


**Figure S10. Biophysical networks shape the gradient organization of functional brain connectivity in the LA5c dataset**. (A) The first two connectivity gradients estimated from the fully connected functional brain network ($FC_{Full}$), and RFNs, i.e., after removing direct connections associated with SC, DC, GC, $GC_d$, RC and $RC_d$. (B) Shift in Gradient 1 and (C) Gradient 2 for different RFNs. Given the complexity of the gradient metric, here to estimate the change, we calculate the Pearson correlation (R) between the gradient map of $FC_{Full}$ and RFNs and then calculate δ as 1-|R| implying dissimilarity in gradient profiles. The gray dots represent the estimated change – dissimilarity – for the null model which was constructed by removing a set of random connections from the functional connectivity network $FC_{Full}$. Note that there was a significant effect of SC on gradients, unlike what was observed in HCP data (Fig. 6). Similarly, while DC and GC emerge as dominant factors impacting Gradient 2, the effect of molecular factors on Gradient 1 is lower than both the physical factors (SC and DC). LA5c included a shorter scan length (~ 5 minutes) than HCP (~ 15 minutes), and some of these differences may reflect that, however, the impact of molecular factors on Gradient 1 seems more nuanced across different robustness tests we performed and requires further testing. Nonetheless, the dominant effect of DC is robustly observed across both the gradients for different methodological and data choices considered in this study. FPN, Frontoparietal Network; DMN, Default Mode Network; DAN, Dorsal Attention Network; LIM, Limbic; VAN, Ventral Attention Network; SOM. Statistical testing assessed the deviation from the null model with *** denoting p-value < 0.001.

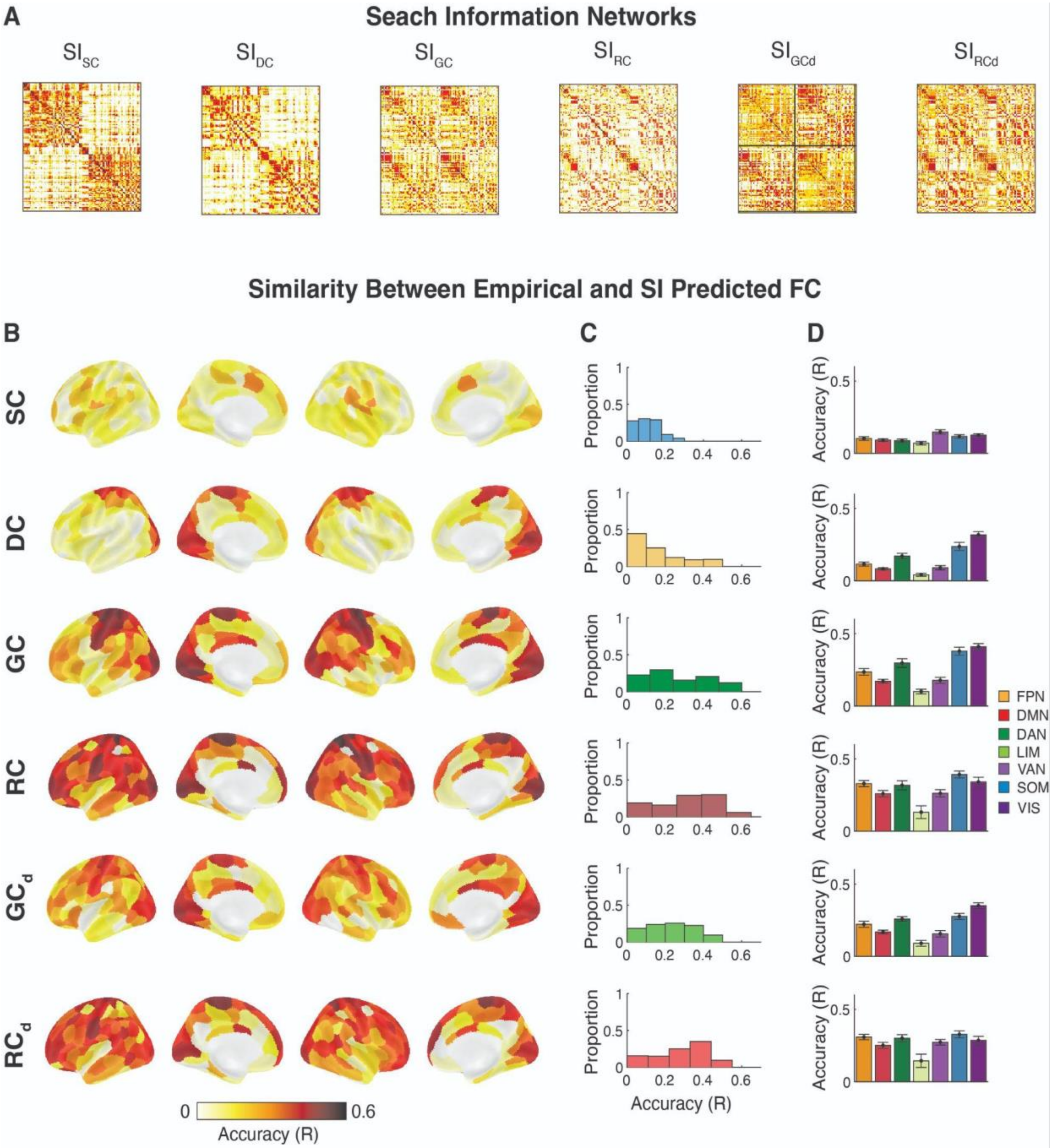


**Figure S11. Modeling the functional connectivity from biophysical networks in the HCP dataset.** (A) Most previous work has focused on modeling on understanding how anatomical wiring constraints and shapes communication[47]. We extend the modeling of FC from biophysical factors using Search Information (SI)[26]. SI measures the amount of information required to traverse the shortest paths between a given pair of brain regions and it has been shown to be a good model for predicting FC. The SI based *predicted* functional connectivity was derived from each of the biophysical networks – SC, DC, GC, $GC_d$, RC and $RC_d$. (B) Accuracy of SI predicted functional connectivity for each brain region. The biophysical networks exhibited differential capabilities in predicting FC (panel B). On average the best predictor of nodewise functional connectivity were RC ($R = 0.31 \pm 0.16$; $P < 0.001$) and $RC_d$ ($R = 0.28 \pm 0.13$; $P < 0.001$). In fact, for 84 out of 200 ROIs (42%), RC was the best predictor of FC.  Accuracy was quantified at the similarity, Pearson correlation (R), of the regional connectivity profile between the SI predicted and empirical functional connectivity ($FC_{Full}$) of each of the 200 brain regions. (C) Distribution of accuracy values among biophysical networks. (D) The average performance within large-scale functional communities. RC and $RC_d$ exhibited similar predictive capabilities across the cortex and large-scale brain networks suggesting neuroreceptor congruence uniformly shapes FC across the cortex.Whereas, the other biophysical networks exhibited localized predictive capabilities. The predictive capabilities of DC, GC and $GC_D$ were primarily constrained to the somatomotor and visual networks (panel D). On the other hand, SC weakly predicted functional connectivity across the brain. FPN, Frontoparietal Network; DMN, Default Mode Network; DAN, Dorsal Attention Network; LIM, Limbic; VAN, Ventral Attention Network; SOM, Somatomotor; VIS, Visual Network.

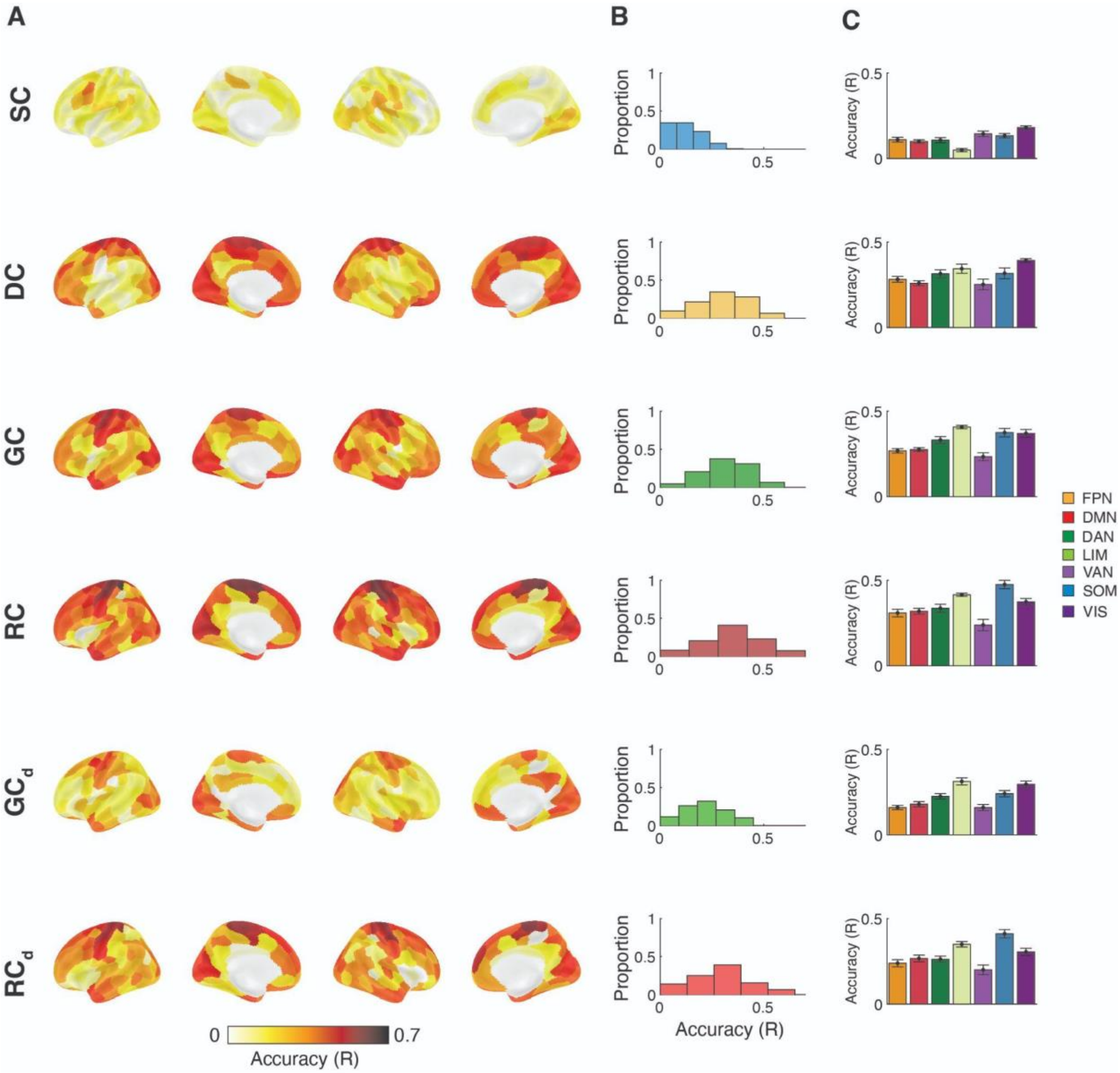


**Figure S12. Modeling the functional connectivity from biophysical networks in the LA5c dataset.** (A) Performance of SI predicted functional connectivity for each brain region. Performance was estimated with a linear model to compare the similarity in the regional connectivity profile between the SI predicted and empirical functional connectivity of each of the 200 brain regions. (B) Distribution of accuracy scored in modeling FC among biophysical networks. (C) The average performance within large-scale functional communities. These findings are aligned with the findings presented in the Fig. S7 for HCP. FPN, Frontoparietal Network; DMN, Default Mode Network; DAN, Dorsal Attention Network; LIM, Limbic; VAN, Ventral Attention Network; SOM, Somatomotor; VIS, Visual Net
